# A weaponized phage suppresses competitors in historical and modern metapopulations of pathogenic bacteria

**DOI:** 10.1101/2023.04.17.536465

**Authors:** Talia Backman, Sergio M. Latorre, Efthymia Symeonidi, Artur Muszyński, Ella Bleak, Lauren Eads, Paulina I. Martinez-Koury, Sarita Som, Aubrey Hawks, Andrew D. Gloss, David M. Belnap, Allison M. Manuel, Adam M. Deutschbauer, Joy Bergelson, Parastoo Azadi, Hernán A. Burbano, Talia L. Karasov

## Abstract

Bacteriophages, the viruses of bacteria, are proposed to drive bacterial population dynamics, yet direct evidence of their impact on natural populations is limited. Here we identified viral sequences in a metapopulation of wild plant-associated *Pseudomonas* spp. genomes. We discovered that the most abundant viral cluster does not encode an intact phage but instead encodes a tailocin - a phage-derived element that bacteria use to kill competitors for interbacterial warfare. Each pathogenic *Pseudomonas* sp. strain carries one of a few distinct tailocin variants, which target variable polysaccharides in the outer membrane of co-occurring pathogenic strains. Analysis of historic herbarium samples from the last 170 years revealed that the same tailocin and receptor variants have persisted in the *Pseudomonas* populations for at least two centuries, suggesting the continued use of a defined set of tailocin haplotypes and receptors. These results indicate that tailocin genetic diversity can be mined to develop targeted “tailocin cocktails” for microbial control.

**One-Sentence Summary:** Bacterial pathogens in a host-associated metapopulation use a repurposed prophage to kill their competitors.

## Introduction

Understanding the factors that influence the spread of pathogens is fundamental for developing disease prevention and treatment strategies. To colonize a plant, bacterial pathogens need to overcome not only the plant’s immune system, but also must compete with surrounding microbes and must defend against bacteriophages. Microbial disease is a frequent outcome in genetically homogeneous agricultural fields, but rarer in genetically diverse wild plant populations (*1*). These wild plant populations are instead typically colonized at low levels by diverse pathogens, with no strain spreading widely (*2*). What prevents the dominance and spread of single lineages as is so often seen in agriculture? The prominent role of host genetics in suppressing pathogenic lineages is well-documented (*3*), but how the surrounding microbiota affect disease dynamics remains largely unknown though of increasing interest (*4*).

Bacteriophages (henceforth phages), bacteria- and archaea-infecting viruses, are the most abundant biological entities on Earth (*5*). Phages can be highly specific in the bacteria they infect, targeting only a few strains of a bacterial species (*6*). The machinery of phages has also been repeatedly co-opted by bacteria to aid in killing competitors (*7*) or in colonizing hosts (*8*). Phages and phage-tail-like elements including bacteriocins or tailocins are hypothesized to be drivers of microbiome composition in host and non-host environments (*9*). To date, the impact of phages and their derived elements on plant microbiota has only been demonstrated in the lab under controlled conditions (*10*). Whether phages and their derived elements are a major suppressor of pathogen strains in the wild remains largely a matter of speculation (*9*) (but see (*11*, *12*)).

We previously found that wild populations of the plant *Arabidopsis thaliana* were colonized by a metapopulation of *Pseudomonas* spp. bacterial pathogens, a genetically-diverse population of *Pseudomonas* populations (*13*). Our prior results suggested that these pathogen populations do not undergo clonal expansions as is frequently observed in agricultural and clinical pathogens. Infections of *Pseudomonas viridiflava* populations, even within an infection of a single plant, consisted of several co-occurring strains and no single strain rising to prominence (*2*). What prevents single pathogenic lineages from spreading? *A. thaliana* immune diversity certainly plays a role in maintaining *Pseudomonas* pathogen diversity; however, our previous work indicated that other members of the plant microbiome also affect the *Pseudomonas* composition (*14–16*). Because phages and their derived elements are common in *Pseudomonas* populations (*17*) we hypothesized that differences in sensitivity to phage components could suppress specific *Pseudomonas* spp. strains in our system.

### The viral cluster VC2 is strongly associated with a pathogenic clade of *P. viridiflava*

To determine whether phages and their derived elements cause turnover in the *Pseudomonas* colonizing *A. thaliana*, we first sought to characterize the abundant phage sequences in a wild *Pseudomonas* metapopulation. Viral sequences integrated within a bacterial genome are the result of lysogeny with a compatible phage (*18*). Identifying compatible lysogeny events can reveal clade-specific phage enrichment. We characterized the presence and evolution of viral elements in 1,524 *Pseudomonas* spp. genomes all collected from *A. thaliana* in Southwestern Germany (Fig. S1) (*19*). More than 85% of these genomes are classified as the ATUE5 clade of *P. viridiflava* (Fig. 1A), an opportunistic pathogen that colonizes *A. thaliana* throughout Europe and the USA (*20*, *21*). We found viral sequences in 99.3% of all the genomes with an average of two viral sequences per genome (Fig. 1B). By using pairwise k-mer distances (*22*) and subsequent k-means clustering, we identified four viral sequence clusters (Fig. 1C).

**Figure 1.**
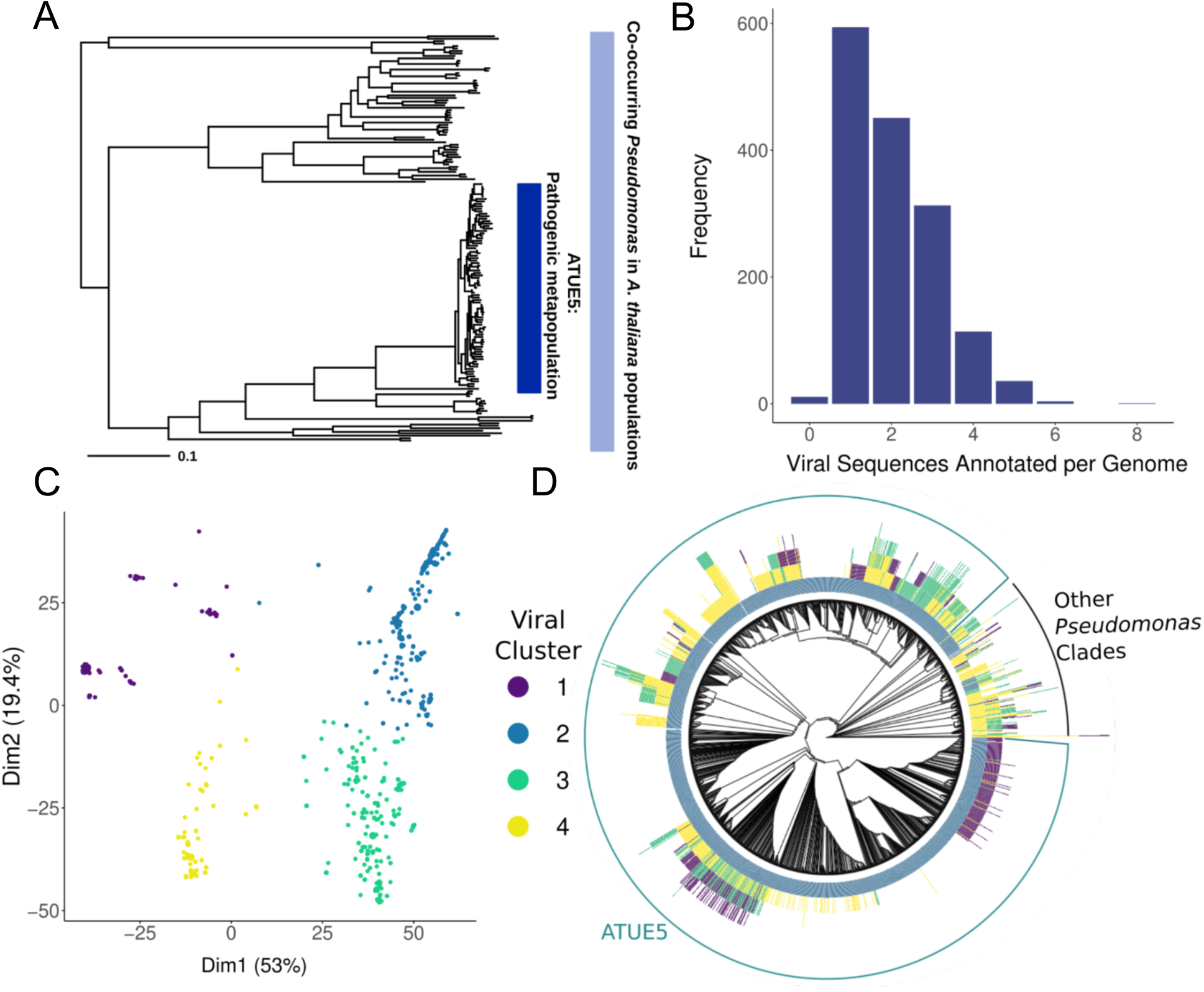
The pathogenic clade of *P. viridiflava* ATUE5 harbors a single highly conserved viral cluster (VC2). **A.** Maximum likelihood concatenated core genome phylogeny of 1,524 pseudomonads that co-occur in the *Arabidopsis thaliana* phyllosphere (light blue bar). The dark blue bar indicates a pathogenic clade of *P. viridiflava* ATUE5 (adapted from Karasov et al. 2018)**. B.** Frequency of viral elements identified in each *Pseudomonas* genome. Each genome harbors between one and eight viral elements. **C.** Principal Component Analysis of viral elements based on pairwise k-mer Mash distances (Ondov et al. 2016) and silhouette analysis reveals four defined clusters. Points are colored as indicated in the inset. **D.** Viral elements of Cluster 2 are present in all genomes of the pathogenic operational taxonomic unit (OTU) ATUE5. The phylogeny shows the relationships among 1,524 *Pseudomonas* strains. Each stacked bar around the phylogeny indicates the presence of a viral element (with a maximum height of eight viral elements). The bars are colored as in C. The outer circle shows strains that belong to ATUE5 (colored in green) and other OTUs (colored in black).

Phylogenetic associations between *Pseudomonas* clades and viral sequence clusters were immediately apparent (Fig. 1D) (*2*), with viral cluster 2 (VC2) found in all ATUE5 pathogenic strains, but less frequent (29%) (Fig. 1D) in non-pathogenic isolates outside of the ATUE5 clade. Furthermore, the gene composition and synteny of VC2 is conserved across ATUE5 strains but has divergent sequences outside the closest relatives to ATUE5 (Fig. 2B). The VC2 sequences in ATUE5 consist of 24 genes that are colinearly integrated in all genomes at the same location — between the *trpE* and *trpG* bacterial genes (*23*, *24*). These VC2 sequences are predicted to encode the structural components typical of a prophage, including baseplates, tail spikes, tubes, sheaths, tail fibers, putative assembly chaperones, and transcriptional regulation proteins.

**Figure 2.**
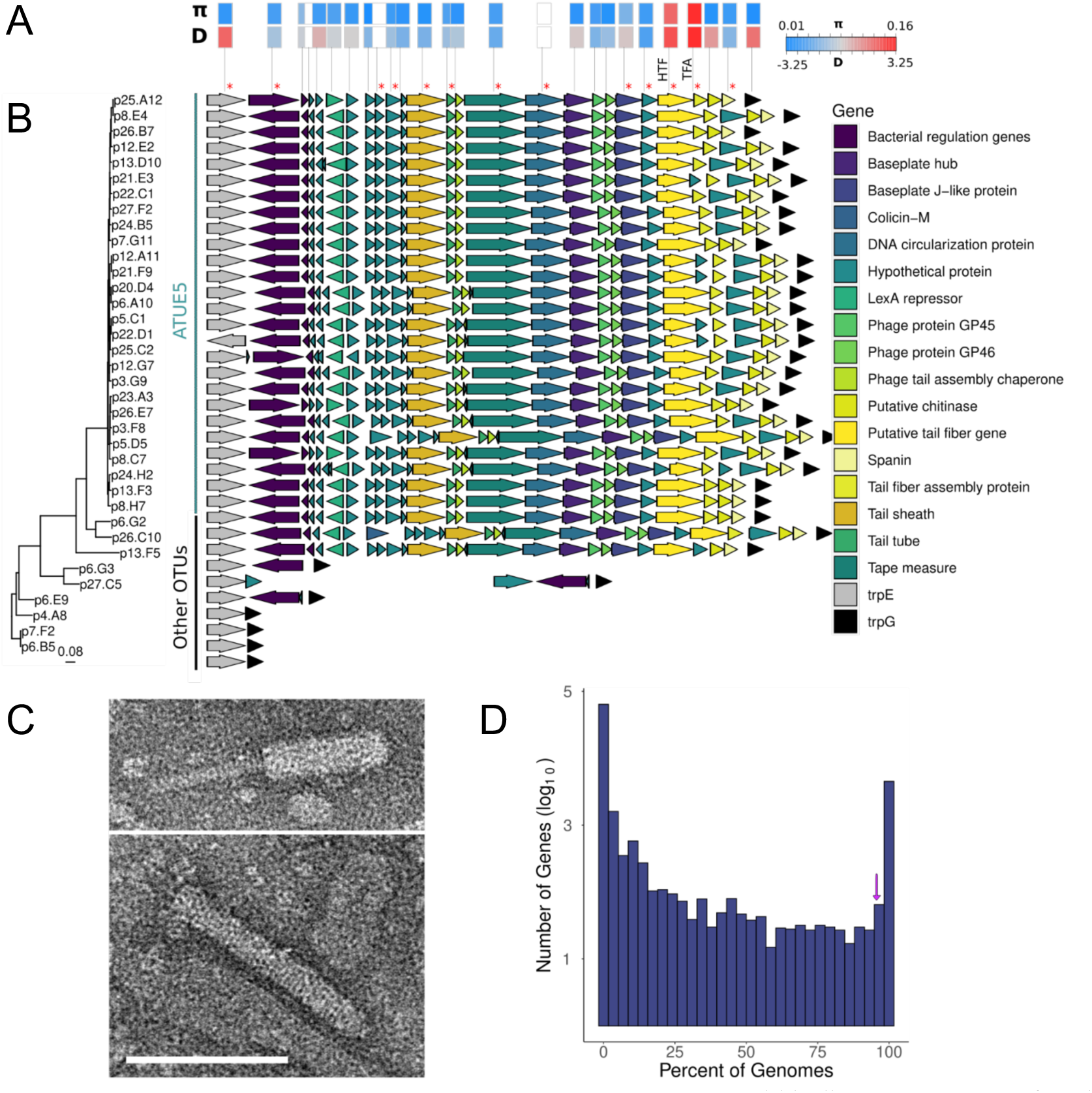
Viral cluster 2 encodes a structurally functional tailocin. **A.** Within-lineage measures of nucleotide diversity (measured as pairwise nucleotide differences (π)) and the site frequency spectrum measured as Tajima’s D (D) for each tailocin gene. White boxes indicate genes that are present in less than 5% of strains (statistics are excluded). Red asterisks (*) indicate proteins found in proteomics analysis. **B.** Genes of Cluster 2 viral elements are syntenic in genomes of ATUE5. The arrows represent tailocin genes and surrounding bacterial genes (gene names are color-coded in the inset). Each row is organized according to its phylogenetic placement. The phylogeny includes 36 *Pseudomonas* representative strains. The vertical lines indicate strains that belong to ATUE5 (colored in green) and other OTUs (colored in black). **C.** Transmission Electron Microscopy (TEM) demonstrates the presence of an assembled tailocin. TEM micrographs show the induced and partially purified tailocin from one representative ATUE5 strain (p25.A12) in both its contracted (top) and (bottom) uncontracted forms. The scale bar, 100nm, applies to all micrographs. **D.** The tailocin is part of the ATUE5 core genome. The histogram shows the frequency of each of the pangenome genes within 1,399 ATUE5 genomes. The eleven most conserved tailocins genes are present in more than 90% of the ATUE5 genomes (marked with a purple arrow in the histogram).

### The viral cluster VC2 encodes a structurally functional tailocin

The analysis of the VC2 gene content revealed that this cluster lacks components necessary for phage replication and encapsulation such as capsid, terminase, integrase, and recombinase proteins, suggesting that the VC2 genomic region does not encode a fully functional phage (*25*, *26*). While the structure of VC2 was not consistent with a functional phage, it did resemble previously described bacteriocins that were derived from phages, dubbed tailocins (*27*). Tailocins are repurposed phages that bacteria use to kill co-occurring bacterial strains (*23*) using the targeting machinery of the phage ancestor.

To test whether the tailocin genomic island produced the predicted tailocin protein complex, we induced tailocin production in a representative strain of ATUE5 (p25.A12) using treatment with mitomycin C (MMC) (*28*), partially purified the lysate, and performed Transmission Electron Microscopy (TEM). Transmission electron micrographs (Fig. 2C, Fig. S2) revealed one rod-like tailocin in the MMC-induced lysates of p25.A12 existing in both the extended (sheath uncontracted) and contracted (sheath contracted, and tube ejected) forms (Fig. 2C). The rod-like structures suggest this tailocin is an R-type contractile tailocin (*29–31*). Image comparison of the two tailocin forms indicated that the tailocins differed in average length: the contracted tailocins were on average 130 nm, while the extended tailocins were on average 144 nm in length. No other intact phage or bacteriocin-like structures were observed in the TEM micrographs.

To validate that the observed tailocin in the TEM image corresponded to our predicted tailocin from VC2, we performed untargeted data dependent LC/MS/MS proteomics analysis (*23*, *32*). Proteomics data confirmed the presence of tailocin proteins annotated in VC2 (Fig. 2B, Fig. S2 and Table S1). No tailocin proteins were found in the sample that was not induced with mitomycin C (MMC) and no other viral proteins or proteinaceous toxins, such as S-bacteriocins, were detected in the tailocin lysates. Together, these results suggest the highly conserved ATUE5 viral sequence is an active and inducible R-type tailocin.

### Co-occurring *Pseudomonas* spp. genomes encode highly diverse tailocin variants

The phylogenetic distribution of the VC2 cluster (Fig. 1, Fig. 2B) suggested that the VC2 genomic island is well-conserved across pathogenic ATUE5 strains. Across the 1,524 *Pseudomonas* spp. genomes analyzed (Fig. 2D) the full complement of the 27 tailocin genes is found in more than 95% of ATUE5 genomes suggesting the tailocin is in the core genome of ATUE5 strains along with essential bacterial genes (*33*). The conservation of the tailocin protein complex across all ATUE5 genomes, its conserved content and its consistent location between *trpE/trpG* (*27*) suggests a single integration of the VC2 tailocin in the common ancestor of ATUE5 that is likely important for fitness in its natural ecological context.

In addition to the conservation of its gene content and collinearity, we found that the VC2 cluster harbors average levels of intra-lineage nucleotide diversity comparable with bona fide *Pseudomonas* core genes (Fig. 2A, Fig. S3A), indicating that the VC2 cluster has likely been evolving under similar levels of selective constraint. However, through calculating nucleotide diversity for each of the VC2 genes individually, we identified greater levels of nucleotide diversity in two tailocin genes, the hypothetical tail fiber (HTF) and the tail fiber assembly (TFA) genes. Moreover, these two genes were also outliers for positive values of Tajima’s D (Fig. 2A, Fig. S3B), a measure of the skewness of the site frequency spectrum, which in this case might indicate gene genealogies with deep divergence times, and the likely existence of distinct haplotypes. It has been previously shown that the HTF and the TFA genes together are key determinants of bacterial host range (*34*), which could explain their high level of diversity and the likely presence of multiple haplotypes.

We formally ascertained 40 and 25 haplotypes for HTF and TFA, respectively. In both cases, the five most common haplotypes accounted for about 80% of the *Pseudomonas* spp. strains. The HTF haplotypes exhibited polymorphism in terms of their sequence length, with the majority falling into four distinct length categories (Fig. 3A). The HTF and TFA haplotypes are significantly associated (Fig. S4, Fisher’s Exact test, p-value = 0.0005). The high positive values of Tajimas’s D suggest that the divergent HTF and TFA haplotypes might have been maintained in the ATUE5 clade for long evolutionary times. Our population genomic analyses suggest that while the tailocin is conserved in presence in pathogenic ATUE5 populations, the different ATUE5 strains encode one of a few divergent tailocin variants, with possible differences in binding and killing activity between the variants.

**Figure 3.**
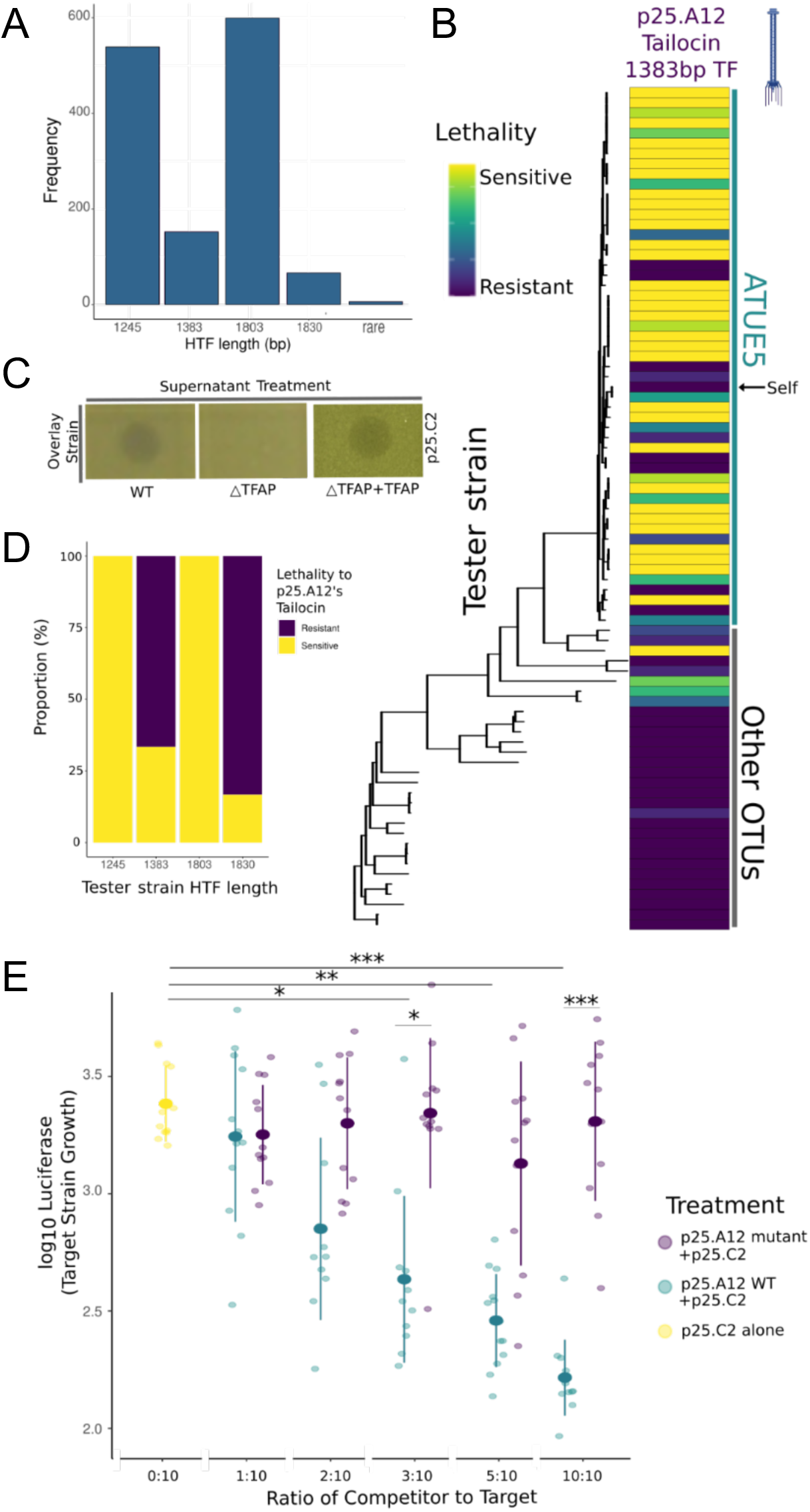
Tailocins target closelyrelated pathogens. **A.** Frequency of HTF nucleotide sequence lengths. There are four highly conserved lengths within the *Pseudomonas* populations. **B.** Tailocins are preferentially used for intra-lineage killing. Soft agar culturesof the *Pseudomonas* strains (rows) were challenged with viral particles extracted from cultures of one strain, p25.A12 (from the 1383bp hypothetical tail fiber length haplotype, column), in 3 technical afor each of 3 biological replicates. The phylogeny includes 83 *Pseudomonas* representative strains and are displayed according to their phylogenetic placement. Vertical lines indicatestrains that belong to ATUE5 (coloredin green) and other OTUs (colored in black). Interactions with the strain’s own tailocin are indicated by the black arrow pointing to self. For each replicate, a strain was given a score of 3 for clear zone of inhibition, a 2 for semi-clear, a 1 for opaque, or a 0 for no killing, and then added together after all 3 replicates. **C.** Knockout of the tail fiber assembly gene disrupts tailocin bactericidal activity. Killing activity is indicated by a clearing in the lawn of the overlay strain. Complementation of the gene on an overexpression plasmid in the knockout strain restores the killing phenotype. **D.** The proportion of tester strains sensitive or resistant to p25.A12’s tailocin, 1383bp hypothetical tail fiber, significantly correlated with the tester strain’s hypothetical tail fiber length haplotype (Fisher’s Exact test, p = 10-8)). **E.** *In planta* coinfections of p25.A12 (competitor) and p25.C2 (target, known to be sensitive to p25.A12). Different ratios of competitor and constant amounts of target strain were used. p25.C2 grown alone as the control. ANOVA test, P-values are shown as p < 0.001 ***, p < 0.01 **, or p < 0.05 *.

### Different tailocin variants kill closely related pathogenic *Pseudomonas*

We hypothesized that like other tailocins, the tailocin variants encoded by the *Pseudomonas* spp. strains are used for interference competition against closely related competitor strains (*23*, *35*). Our system provides a unique opportunity to characterize killing activity of tailocins produced by co-occurring microbes in the same wild plant populations. The relatedness of these microbes spans the spectrum from closely related (ATUE5, <1% genome wide sequence divergence) to distant commensals (other ATUEs, >20% genome wide sequence divergence).

We hypothesized the ATUE5 tailocin is used for interference competition either against (i) other pathogenic pseudomonads that also produce a tailocin, (ii) commensal pseudomonads that mostly do not produce the VC2 tailocin, or (iii) other bacterial strains in the phyllosphere community. To test this, the tailocins from 3 strains and 2 representative HTF length haplotypes were induced and partially purified, then concentrated tailocin lysates were applied to 55 ATUE5 strains, 28 other ATUE strains, and over 50 other bacterial strains isolated from the phyllosphere (taxonomy unknown). We found that, as expected, ATUE5 strains were resistant to their own produced tailocin (*23*). The killing assay also revealed that non-ATUE5 *Pseudomonas* strains were sensitive in only 6% of ATUE5-derived tailocin treatments, while 40% of ATUE5 treatments with ATUE5-derived tailocins exhibited sensitivity (Fig. 3B, Fig. S6). To test whether other non-*Pseudomonas* community members were sensitive to the tailocin treatments, we tested fifty colonies from the surrounding phyllosphere, and found that none were sensitive to tailocin treatment. An agriculturally relevant *Pseudomonas* strain, DC3000, was also tested and found to be sensitive to one of the tested tailocins (p25.A12). In conclusion, the different tailocin variants appear to exhibit partial killing specificity to different subsets of other ATUE5 pathogenic *Pseudomonas*, only 6% of the other *Pseudomonas* commensals, and not to other phyllosphere strains except for the above mentioned DC3000 case.

To verify that the tailocin is responsible for the observed killing activity, we generated a TFA-deficient mutant. Tailocins collected from the mutant (p25.A12ΔTFA) lost killing activity against all strains tested (Fig. 3C, Fig. S5), and the killing phenotype could be restored with an overexpression plasmid containing the TFA gene. These results suggest the tailocin is necessary for the observed killing activity.

To determine whether the TFA or HTF haplotypes are associated with killing activity, we analyzed whether the tailocin killing spectrum correlates with a bacterial tester strain’s TFA or HTF haplotype. Rather than a correlation, we found an array of different killing spectra, with the tail TFA/HTF haplotypes tested all displaying broad killing against other ATUE5 strains (Fig. 3B and Fig. S6). Although, we found there to be no significant associations between killing spectra and TFA or HTF haplotype, we did identify a significant association between the length of the HTF and its killing spectrum (Fig. 3A and 3D, Fisher’s Exact Test p = 10^-8^.). This suggests that tail fiber sequence length could be important in binding to target cells and is likely associated with an outer membrane receptor gene.

We next aimed to determine whether tailocin killing is observed in the plant host. We performed competition assays *in planta*. Strain p25.A12’s tailocin was able to kill a particularly pathogenic strain, p25.C2, in all three killing assays (Fig. 3B, 3rd row). To test whether the target strains would be killed by p25.A12, we grew strains together using a variant of p25.C2 tagged with luciferase to measure its growth. As a control, we also competed p25.C2 with p25.A12ΔTFA. Five ratios of p25.A12 and p25.C2 were used to determine whether the concentration of the tailocin strain affected killing, while controlling the abundance of the target strain. We found that p25.A12 significantly reduced the growth of p25.C2 in the three highest ratios tested (Fig. 3E). The 10:10 and 3:10 ratios significantly differed from the p25.A12ΔTFA competitive assay, suggesting that having a functional tailocin suppresses growth of p25.C2 and that p25.A12 is no longer able to significantly reduce the growth of p25.C2 when lacking the tailocin. We found that p25.C2 grown with p25.A12ΔTFA was never significantly different than p25.C2 grown alone, suggesting that the tailocin is an effective competitive mechanism in the plant host. These results were replicated in competition assays done in test tubes (Fig. S7), however in these trials p25.A12ΔTFA competing with p25.C2 significantly differed from the control for the highest ratio. This result could indicate that p25.A12 utilizes another competitive mechanism in the absence of tailocin, which proves ineffective at lower starting abundances. Together, these results suggest that tailocin killing is important in killing closely related competitors *in planta*.

### The tail fiber is co-evolving with its O-antigen receptor in natural populations

The differences we observed between strains in their susceptibility to the tailocin (Fig. 3B) indicated that these strains may encode an unidentified receptor that has different variants. To identify the receptor, we conducted a transposon mutagenesis screen (*36*) in which we screened a mutant library of a susceptible strain (p25.C2) for insertions that render the strain resistant to the p25.A12 tailocin (Fig. 4A). Insertions (putative knockouts) in seventy genes were significantly associated with resistance to the tailocin. Six of these genes lie within an O-antigen biosynthesis gene cluster, a locus previously implicated in resistance to phage and tailocin tail fibers (*37*). These results suggested that differences between strains in their O-antigen may explain differences in tailocin susceptibility. Comparative DOC-PAGE analysis of the lipopolysaccharide (LPS) isolated from two resistant and two susceptible strains showed the resistant strains lack a high molecular weight polysaccharide chain that is present in the susceptible strains. Subsequent chromatography-mass spectrometry (GC-MS) (Fig. 4C) revealed the susceptible strains had relatively high amounts of rhamnose compared to the resistant strains (Table S2). Our finding of the higher content of rhamnose, along with the expanded size of the LPS in tailocin resistant strains, agrees with the previous studies suggesting the affected gene locus might be involved in the biosynthesis of the O-antigen in *Pseudomonas* (*38–40*). Here, we propose the O-antigen is likely a receptor for the focal tailocin.

**Figure 4.**
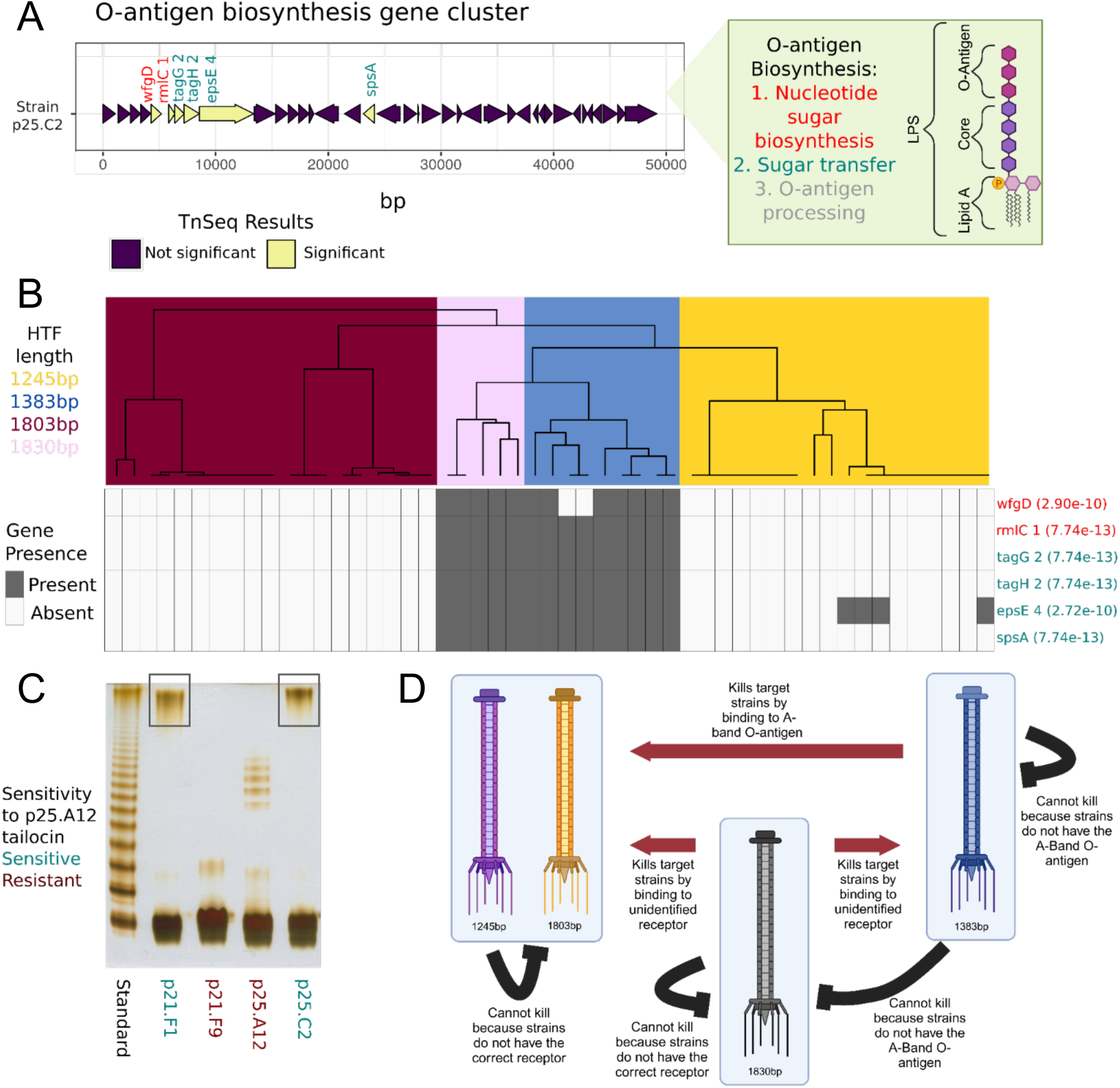
The O-antigen is important for tailocin lethality and is co-evolving with the tailocin HTFP. **A.** Gene plot of the a previously characterized O-antigen gene cluster and found in ATUE5 strains. Six of the seventy significant TnSeq genes in this study were found in this gene cluster, and are shown in yellow. Gene names are colored by the step in O-antigen biosynthesis pathway. **B.** The top dendrogram is from hierarchical clustering of the *PanKmer* output for the hypothetical tail fiber gene, colored by length. Rows represent the six significant O-antigen genes. Grey corresponds to gene presence and white to gene absence. Fisher’s exact test *p*-values between genes and length-based clusters are shown in parenthesis **C.** DOC-PAGE profile of silver stained LPS (1 µg each lane) isolated from a subset of ATUE5 strains. The standard is of Salmonella enterica Ser. Typhimurium (S-type LPS). Grey boxes indicate high molecular weight O-chain. **D.** Our working model of tailocin killing and lethality in *P. viridiflava* isolates. Red arrows indicate known significant patterns of killing. Black lines show known significant patterns of non-killing. Figure was created with BioRender.com.

Given that strains are resistant to their own tailocins (*41*), we posited that the tailocin and receptor loci should co-evolve. To test this, we measured congruence between tail fiber and LPS haplotypes (Fig. 4B) and found the tail fiber and LPS haplotypes to statistically co-vary (Fisher’s exact test p-values < 2.72e-10 for every association between presence/absence of the gene and the length-based clusters; Fig. 4B). Knowing the tail fiber haplotype allows for prediction of the LPS haplotype. In total, these experiments support a model in which the tail fiber and LPS of co-occurring *Pseudomonas* strains are evolving in concert in wild *A. thaliana* populations (Fig. 4D).

### The divergent tailocin and LPS haplotypes have been maintained across time and space

To ascertain the timescales of the emergence of the tailocin and different tailocin variants, we sought to retrieve *Pseudomonas* genomes from historical *A. thaliana* herbarium specimens collected in the last two centuries. Only historical *Pseudomonas* genomes can reveal whether extant populations are the result of population continuity or population replacement. We employed state-of-the-art ancient DNA retrieval and sequencing techniques (*42*) to sequence leaves of 35 *A. thaliana* herbarium specimens using shot-gun genomics (Table S3). We mapped the herbarium-derived reads to the ATUE5 reference genome strain to identify *Pseudomonas* reads and found seven samples with more than 60% of the genome covered by at least one sequencing read (mean genome coverage 0.97-8.7x) (Fig. S8, Table S3). The *Pseudomonas*-derived reads showed patterns of DNA damage and fragmentation typical of ancient DNA (*43*), which authenticated their historical origin (Fig. S9). Our analysis revealed that herbarium specimens preserve the genomes of highly abundant bacterial genera of the *A. thaliana* phyllosphere.

To place the historical *Pseudomonas* genomes in the context of extant genetic diversity, we first used patterns of homozygosity support to identify herbarium specimens most likely to be colonized by a single dominating *Pseudomonas* spp. strain (Fig. S10). This analysis allowed us to select three historical samples (mean genome depth of 8.2 - 9.6x), which we combined with 65 present-day *Pseudomonas* genomes to build a phylogenetic tree using whole-genome Single Nucleotide Polymorphisms (SNPs). The three historical samples were placed within the present-day ATUE5 diversity (Fig. 5), showing the genetic continuity of this clade for at least the last 177 years. Moreover, the mappings of the historical samples against the ATUE5 reference genome identified the presence of the tailocin in all three historical bacterial genomes (Fig. 5A). In order to ascertain the tailocin variants encoded in the historical genomes and to assess whether the gene colinearity and the insertion location in the bacterial genome are conserved in the historical genomes, we carried out *de novo* assembly of historical *Pseudomonas*-derived reads. We obtained an average of 9 contigs per sample (average sequence length of ∼3.4 Kb) that were sufficient to show that both the insertion place of the tailocin and the colinearity of its genes are conserved in the historical genomes (Fig. 5A). Moreover, we ascertained that each historical genome carries a different tailocin variant and that all historic variants are segregating in present-day populations (Fig. 5B, Fig. S11A). The polyphyletic distribution of the HTF and TFA gene haplotypes (Fig. 5B, Fig. S11A) is likely the result of recombination at this locus (*34*).

**Figure 5.**
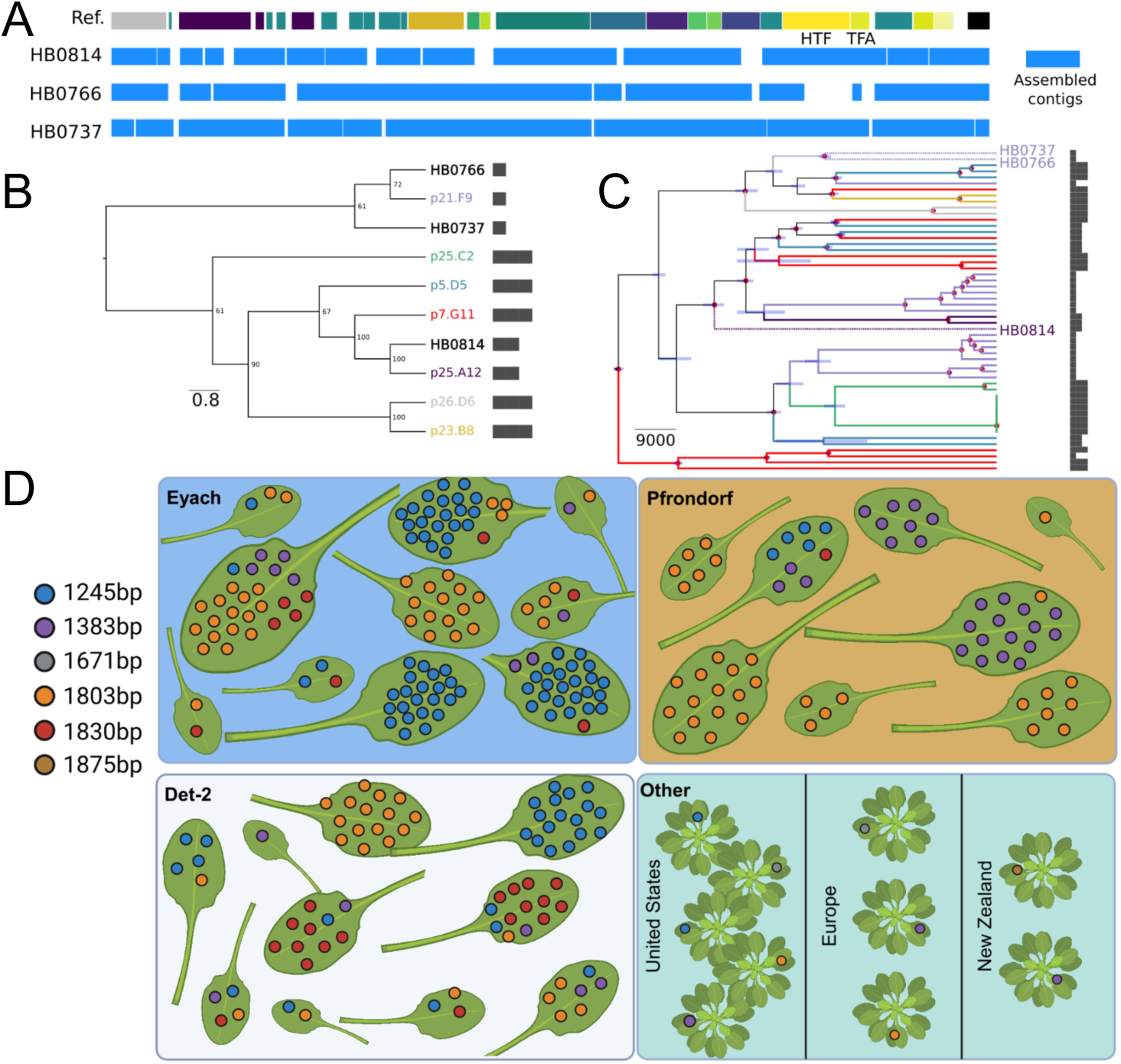
The same length haplotypes of the tail fiber (HTF) are present in contemporary and century-old historical samples and the HTF haplotype diversity is maintained at the leaf scale and broader geographical scales. **A.** *De-novo* assembled contigs spanning the tailocin genomic region for the historical samples. The reference track on the top represents the genes present in the ATUE5 reference genome. The blue boxes depict the *de-novo* assembled contigs. **B.** Maximum-Likelihood Neighbor-Joining tree of the hypothetical tail fiber gene translated sequence. Historical samples (HB) are placed in the context of the most common haplotypes. HTF lengths are shown with bars. **C.** Bayesian tip-date calibrated phylogeny representing the evolutionary relationship between historical and modern *Pseudomonas* sp. strains. The tips and branches are colored based on the hypothetical tail fiber gene haplotype. Only historical samples labels are shown with stars. The node bars represent the 95% Highest Posterior Density Intervals of the estimated time and the nodes marked with red dots represent those with posterior probability of 1. HTF lengths are shown with bars. **D.** A subset of eight to ten plants from the three German populations (blue, orange, and white panels), and 10 Pseudomonas syringae isolates collected from plants globally, downloaded from NCBI (green). Each leaf is a leaf from a single plant. Colored dots indicate a single bacterial isolate found and its corresponding HTF length. Plants in the bottom right panel indicate plants not found within the same population. Figure was created with BioRender.com.

We used our phylogenetic reconstruction to determine that the tail fiber assembly gene haplotypes of the historical genomes are closely related to different modern haplotypes (Fig. 5C, Fig. S11B), suggesting that the same haplotypes have circulated in the *Pseudomonas* metapopulation for the past 177 years.

The observed persistence of HTF length haplotypes within the bacterial metapopulation may result from a potential artifact introduced by population structure at the plant, plant patch, or site level. At these levels, specific haplotypes could reach fixation, creating the appearance of the maintenance of multiple haplotypes when analyzing all populations collectively. To rule out this possibility, we looked at the co-occurence of the HTF length haplotypes at the level of single leaves, all collected within a single site in the Germany population as well as at broader geographic locations through analysis of 10 *Pseudomonas syringae* genomes from NCBI (5 isolates collected from United States, 3 from Europe, and 2 from New Zealand). We found that HTF lengths were co-occurred in single leaves and in all three German populations (Fig. 5D). Additionally, we found that the 1245bp, 1383bp, and 1803bp HTF lengths in other countries, in addition to two other lengths. These results suggest that HTF lengths are conserved and maintained at the microscale and over great geographic distances.

## Discussion

Our results have two major implications. First, we discovered that a tailocin, a repurposed phage, is conserved as part of the core genome of metapopulations of the pathogen *P. viridiflava*. Its conservation in presence for at least tens of thousands of years and the sequence conservation of its genes (other than the TFA/HTF) indicate that the tailocin is likely essential (or nearly so) for survival in *Pseudomonas* pathogen metapopulations. This finding is consistent with previous work that suggests that antimicrobial nanomachines such as the Type Six Secretion System are important for survival in the plant environment (*44*). Despite its conservation in presence, multiple allelic variants of the tailocin have been maintained in these populations. We found that the allelic variants target different strains of related pathogenic *Pseudomonas* species/subspecies via differences in the LPS enriched outer leaflet of the outer membrane (*23*). Similar to the coevolution of toxin-antitoxin systems, the tail fiber and assembly proteins of an ATUE5 strain are coevolving with the LPS membrane composition (*45*). The strong evolutionary conservation and correlation between loci suggests that inter-*Pseudomona*s-strain competition via the tailocin is a strong and persistent selective pressure on *Pseudomonas* pathogens.

Second, our results revealed that the same tail fiber/LPS variants have been maintained in the *Pseudomonas* populations for the past two hundred years. Bacteria are capable of rapid adaptation (*46*). The fact that a defined set of variants is maintained in these populations suggests selective constraints on the evolution of this system, consistent with what has been found in viral evolution (*47*). The presence of a limited set of tail fiber haplotypes in the metapopulation could reflect a limited panel of resistance mechanisms.

Together, these findings provide a roadmap for identifying tail fiber specificities to different strains and the possibility of determining the mechanism of this specificity. Tailocin therapy, like phage therapy, has been proposed as a possible alternative to small molecule antibiotics (*48–50*). Proof-of-concept studies demonstrated that tailocins can be used to suppress specific pathogens in animal (*48*) and plant (*49*) models. With any antimicrobial treatment, there is a risk that the target bacterium will evolve resistance. Our findings indicate that, considering the probable constrained array of resistance mechanisms, the tailocin genetic diversity of a pathogen metapopulation can be mined to discover “tailocin cocktails”. These cocktails have the potential to target the metapopulation in parallel, thereby reducing the likelihood of resistance evolution. In the future, the mining and characterization of the tailocin repertoire from diverse wild *Pseudomonas* populations will unveil the possible combinatorics between the tail fiber and LPS composition, as well as the timescales in natural settings over which tailocin resistance emerges.

## Acknowledgements

We thank Sophien Kamoun, Patricia Lang, Gal Ofir, Dmitri Petrov and Detlef Weigel for comments that improved this manuscript.

## Funding

This study was supported by startup funds from the University of Utah and by NIH grant R35 GM150722-01 (TLK), The Leverhulme Trust (Philip Leverhulme Prize) (HAB), the Royal Society (RSWF\R1\191011) (HAB), and an equipment grant from the UK Biological Sciences Research Council - BBSRC (BB/R01356X/1) to University College London. Proteomics mass spectrometry analysis was performed at the Mass Spectrometry and Proteomics Core Facility at the University of Utah. Mass spectrometry equipment was obtained through a Shared Instrumentation Grant #S10 OD018210 01A1. This work was supported in part by the U.S. Department of Energy, Office of Science, Basic Energy Sciences, Chemical Sciences, Geosciences and Biosciences Division, under award #DE-SC0015662 (PA) at DOE Center for Plant and Microbial Complex Carbohydrates at the CCRC ; and by NIH grant R24GM137782 to (PA) at the National Glycoscience Resource-CCRC Service and Training.

## Author contributions

Biological discovery and conceptualization: TB, TLK, HAB. Methodology: TB, TLK, HAB, SL, ADG, JB. Experiments: TB, DB, AMM, SS, LE. Computation: TB, SL. Writing: TB, TLK, HAB, SL, JB.

## Competing interests

The authors declare that they have no competing interests.

## Data and materials availability

All data and code are available in the supplementary materials or at the git pages: https://github.com/talia-backman/Ps1524_tailocin https://github.com/smlatorreo/Pseudomonas_tailocin.

## Materials and Methods

### Bacterial strains, plasmids and growth conditions

The strains used in this study are detailed in Table S2. Bacteria were grown on nutrient lysogeny broth agar (LB agar) and in nutrient LB media at 28°C for *Pseudomonas* spp. strains and 37°C for *Escherichia coli* strains. All liquid cultures were incubated with shaking at 180 rpm. Growth media were supplemented with kanamycin (50 mg mL−1, Km50), nitrofurantoin (100 mg mL -1, NFN100), or tetracycline (20 mg mL -1, TET20). All strains were stored at −80°C in 20% glycerol [v/v]. For plasmid preparation, *E. coli* transformants were cultured in LB. TET20 was used to select *E. coli* transformants. LB supplemented with sucrose, and nitrofurantoin (10 g/L tryptone, 5 g/L yeast extract, 10% (W/V) filtered sucrose, 15 g/L agar, NFN100) plates were used in the selection of the second *P. syringae* homologous recombination event.

### Viral Identification and Clustering

*VIBRANT* v1.2.1 (*19*) was run on the 1,524 assembled *Pseudomonas* genomes and 10 NCBI genomes to identify viral sequences. To group the 3,104 annotated viral sequences from *VIBRANT* into genetically similar clusters the pairwise mutation distance was calculated between all 3,104 viral sequences with *Mash* v2.2.2 (*22*), a MiniHash-based technique that reduces sequences to compressed k-mer representations (sketches). *Mash* was run ignoring single copy k-mers that are more likely the result of sequencing errors, and a sketch size of 10,000 which corresponds to the number of non-redundant min-hashes that are kept for the pairwise distance calculation. This resulted in a 3,104 x 3,104 pairwise mutation distance matrix. Next, the viral sequences were clustered from the pairwise mutation distance matrix with principal component analysis and K-means clustering (*51*, *52*). Silhouette analysis (*53*) was performed to determine the optimal number of clusters for the data.

### Annotation of the viral cluster 2 (VC2) gene cluster

A highly conserved viral sequence, the VC2 gene cluster, was identified in assembled genomes using VIBRANT v1.2.1 and groups of orthologous genes (orthogroups) were identified using *panX* (*54*). Insertion sites for VC2 gene clusters were identified by searching for orthologs of known viral sequence flanking genes: *mutS*, *cinA*, *trpE*, and *trpG* genes. VC2 was always found between *trpE* and *trpG*. Functional gene annotations were determined by orthologous viral genes previously characterized in *VIBRANT* and *BLAST*. Tailocin gene clusters were visualized with the R package “gggenes”.

### Viral particle extraction and partial purification

Overnight cultures were back-diluted into 50 mL fresh LB to extract and isolate tailocins from the various *Pseudomonas* spp. strains. When cultures reached an exponential growth phase (an OD 600nm of 0.4-0.6) 5 µg mL−1 of mitomycin C (MMC, Selleck Catalog No. S8146) was added to induce the bacterial SOS response and tailocin induction. The cultures were then incubated at 28°C for a minimum of 18 hours. Cultures were then centrifuged at 4°C for 1 hour at 1,400g to pellet cell debris. The supernatants were sterilized by filtration with 0.2 μm cellulose acetate filters. To precipitate viral particles, 40% [w/v] of ammonium sulfate was slowly added to the filter-sterilized tailocin lysate while stirring on ice. The lysate was left stirring on ice for at least 18 hrs. After 18 hrs at 4°C, the ammonium sulfate was pelleted by centrifugation at 2,090g for 2 hrs at 4°C. The supernatant was discarded, and the pellet was resuspended in 500 μL cold P-buffer (100 mM NaCl, 8 mM MgSO4, 50 mM Tris-HCl, pH 7.5) and stored at 4°C. Tailocin lysates were used fresh or for up to two weeks.

### Visualization of viral particles using transmission electron microscopy

One negatively stained specimen (from strain p25.A12) was prepared and imaged in the following manner: 3.5 μl of sample was placed on a carbon-coated, transmission EM grid that had been glow-discharged to make it more hydrophilic. After approximately one minute, filter paper was used to remove excess solution from the grid by touching the side of the grid. This was followed by two steps of brief (approx. 1-2 seconds) wash in P-buffer and blotting with filter paper in the same manner. Next, this process was repeated twice but with the negative stain solution (1% ammonium molybdate). Finally, the grid was incubated for 15-20 seconds in a droplet of 1% ammonium molybdate, blotted again, and allowed to air dry. Negatively stained specimens were viewed on a ThermoFisher Tecnai 12 transmission electron microscope operated at 120 kV. Images were recorded on a Gatan UltraScan camera.

### Protein mass spectrometry

Tailocins partially purified as described above were treated with or without MMC and resuspended in 50 µM triethylammonium bicarbonate and 5% SDS (1x S-TRAP buffer) and frozen at -80°C until digestion. Total protein content was quantified using the Pierce™ BCA Protein Assay Kit™ (ThermoFisher Scientific, Waltham, MA) and 10 µg of protein was diluted into 25 µL of 1x S-TRAP buffer. The protein was reduced with 20 mM dithiothreitol at 37°C for 30 minutes and alkylated with 40 mM iodoacetamide at room temperature for 45 minutes in the dark. Samples were acidified with phosphoric acid and digested using micro-S-TRAP™ columns (Protifi, Fairport, NY) with 0.5 µg of trypsin/Lys C per sample according to the manufacturer’s instructions. The peptides were dried to completion, resuspended in 300 µL of 0.1% trifluoroacetic acid and desalted using Pierce™ Peptide Desalting Spin Columns (ThermoFisher Scientific, Waltham, MA) according to the manufacturer’s instructions. The peptides were resuspended in 40 µL of 0.1% formic acid for LC-MS/MS analysis.

### Mass spectrometry analysis of samples

Reversed-phase nano-LC-MS/MS was performed on an Dionex UltiMate 3000 RSLCnano system coupled to a ThermoFisher Scientific Q Exactive-HF orbitrap mass spectrometer equipped with a nanoelectrospray source. 1 μg of each sample were first trapped on a 2 cm Acclaim PepMap-100 column (ThermoFisher Scientific, Waltham, MA) with 5% acetonitrile at 5 μl/min and at 5 minutes the sample was injected onto the liquid chromatograph reverse-phase Acclaim™ PepMap™ 100 C18 2.0 µm nanocolumn (ThermoFisher Scientific, Waltham, MA). A 500 mm long/ 0.075 mm inner diameter nanocolumn heated to 35°C was employed for chromatographic separation. The peptides were eluted with a gradient of reversed-phase buffers (Buffer A: 0.1% formic acid in water; Buffer B: 0.1% formic acid in 100% acetonitrile) at a flow rate of 0.2 µL/min. The LC run lasted for 85 minutes with a starting concentration of 5% buffer B increasing to 28% buffer B over 75 minutes, up to 40% buffer B over 10 minutes and held at 90% B for 10 minutes. The column is allowed to equilibrate at 5% buffer B for 20 minutes before starting the next data acquisition. The Q Exactive-HF mass spectrometer was operated in data-dependent acquisition MS/MS analysis mode selecting the top 20 most abundant precursor ions between 375-1650 m/z at 60,000 resolution for fragmentation at 15,000 resolution.

### Mass spectrometry data analysis

The raw data was analyzed using Proteome Discoverer 3.0 software with the SEQUEST algorithm against the uniprot_ref_pseudomonas_viridiflava database (1-18-2023 version with 4,389 proteins) or p25.A12 database. An allowance was made for 2 missed cleavages following trypsin/Lys C digestion. No fixed modifications were considered. The variable modifications of methionine oxidation and cysteine carbamidomethylation were considered with a mass tolerance of 15 ppm for precursor ions and a mass tolerance of 0.02 Da for fragment ions. The results were filtered with a false discovery rate of 0.01. A minimum of 1 unique peptide should be reported for all proteins identified.

### Allelic diversity and nucleotide sequence data analysis

Orthogroups that were previously identified using *panX* were included in the analysis. The nucleotide and peptide sequences were aligned with *Clustal Omega* (*55*), and then a codon-based nucleic acid alignment was generated with *pal2nal* (*56*). This is critical to ensure the nucleic acid alignment is aligned codon-by-codon, to know whether substitutions result in a synonymous or nonsynonymous amino acid change. The output aligned file was saved in fasta format and used as an input file for population summary statistic calculations in R using the package “PopGenome” v2.7.5 (*57*). The level of sequence diversity among ortho-groups was compared using summary statistics ‘π’ (average pairwise difference or average number of nucleotide diversity per site) (*58*), ‘θ’ (population mutation parameter or number of segregating sites) (*59*), whereas Tajima’s D (*60*) was used as a proxy for the site frequency spectrum (SFS) and thus to determine the demographic and selective forces shaping the SFS of tailocin genes.

### Ascertainment of tailocin haplotypes

To determine how many haplotypes of tail fiber assembly and hypothetical tail fiber genes there are in the pathogenic strains in our study, nucleotide and peptide alignments from *Clustal Omega* (*55*) were analyzed using the R package *pegas* (*61*). We used the haplotype function to determine how many unique haplotypes are in the ATUE5 strains. The most common haplotypes were used for downstream analyses.

### Mapping of reads from historical samples to ATUE5

A set of 35 herbarium *Arabidopsis thaliana* samples collected between 1817 - 1957 from Southern Germany (Table S3) (*62*) were screened for *Pseudomonas* sp. presence. Adapter sequences from the raw reads were trimmed and merged using *Adapterremoval* V.2 (*63*) and subsequently mapped to the *Pseudomonas* p25.A12 assembly using *bwa aln* v.0.7.17 (*64*) with seed deactivation to allow the alignment of of substitutions present at the termini of the reads, *e.g.* those arising from DNA deamination in historic material (*42*). A total of 7 samples for which the *Pseudomonas* reference genome was covered in at least 60% were kept (Table S2). To authenticate the historic origin of the bacterial reads, the proportion of C-to- T (or G-to-A) substitutions as well as the fragment size distribution of the reads were computed using mapDamage v.2.2.1 (*65*) (Fig. S8). We analyzed patterns of homozygosity in the mapped reads to keep only samples which were likely dominated by single *Pseudomonas* sp. lineages (Fig. S10). As a result, only the samples HB0737, HB0766 and HB0814 were kept for downstream analyses.

### Ascertainment of tailocin variants in historical genomes

In order to identify reads aligning to the whole spectrum of tailocin genetic diversity, different haplotypes of the highly polymorphic tail fiber assembly and hypothetical tail fiber genes (Fig. 3A) were added to the p25.A12 reference genome, before the historical samples were mapped as described above. Reads covering the Tailocin region surrounded by the flanking genes *trpE* and *trpG* as well as all reads covering any haplotype of the mentioned genes were subset from the raw reads and used as input for *de-novo* assembly using *SPAdes* v.3.13.0 (*66*). Samples HB0737, HB0766 and HB0814 yielded 10, 9 and 14 assembled contigs, respectively (Table S4). To find collinearity between the *de-novo* assemblies and the reference Tailocin region, we used *minimap* v.2.1 (*67*). Aminoacid translated sequences of the historical isolate-specific haplotypes for the tail fiber assembly protein and Putative tail fiber protein were merged together with the haplotypes segregating in the observed diversity. Finally, we used *Clustal Omega* v.1.2.4 (*68*) to generate a multiple alignment and classify the haplotypes in the historical samples.

### Whole-genome phylogenetic reconstruction combined present-day and historical genomes

A set consisting of 50 ATUE5 modern and 3 historic *Pseudomonas* spp. strains were used to create a phylogenetic reconstruction. We used the mapped reads to perform individual *de-novo* variant callings using *bcftools mpileup* and *bcftools call* v1.11 (*69*) removing those reads with mapping quality values lower than 30. We then merged all individual calls using *bcftools merge* v.1.11 (*69*) and removed those samples with less than 60% of the called variants. We finally kept only biallelic SNPs and sites with a maximum missing information of 10% of the remaining samples. The number of retrieved variants were 176,528 SNPs segregating among 53 strains (50 modern and 3 historic).

Using the genomic SNPs we sought to reconstruct the phylogenetic relations among the 53 strains. We used *IQ-TREE* V2.2.0.3 ((*70*)) in combination with *ModelFinder* ((*71*)), and ultrafast bootstrap *UFBoot* ((*72*)) to estimate a Maximum-Likelihood phylogenetic tree. Variant sites that are likely to arise due to putative recombination were identified using *ClonalFrameML* v.1.12 ((*73*)) and the output was used to mask putative recombinant positions in the original genomic SNP array. We finally used *BEAST2* (*74*) to jointly estimate the phylogenetic relations among the strains as well as the relative time of emergence for all the tree nodes. For this purpose we used the collection dates (Table S3) and a previously estimated evolutionary rate ((*75*)) as priors. In order to minimize parameter estimation we chose HYK as the evolutionary model and to avoid demographic history assumptions we used a Coalescent Extended Bayesian Skyline approach(*76*).

### Testing bacterial sensitivity to tailocins with spot test phenotypic assays

To test the sensitivity of the different bacterial strains to the tailocins, soft agar assays were performed using an adaptation of a protocol from Vacheron et al. 2021 (*35*). 750 μL overnight cultures of each strain was mixed with 25 mL of LB soft agar (0.8%). The mixture was poured into a square plate and left to harden. Then, aliquots of 3-5 μL of concentrated viral particles suspension were applied to the agar, along with a control of P-buffer alone and non-induced cultures in serial dilutions. The plates were incubated overnight at 30°C. Bacterial sensitivity or resistance to the viral particles was assessed after 24 hours. When testing serially diluted tailocins, no plaques were observed, suggesting the killing agent is nonreplicative and not a phage. When testing the uninduced tailocin control samples (not induced with mitomycin C), if lethality was observed, the interaction was deemed to be inconclusive. Spot tests were performed in three biological and three technical triplicates.

### Testing bacterial sensitivity to tailocins in culture

To test the sensitivity of a known sensitive strain (p25.C2) to the mutant tailocin from 25.A12△TFA, p25.C2 was grown overnight then diluted by a factor of 1:10 in fresh LB medium. Once the culture reached growth phase, cultures were supplemented with 5% p25.A12 tailocin, 5% p25.A12△TFA tailocin, or a buffer control. OD measurements at a wavelength of 600 nm were measured in a 96-well plate using a microplate reader (TECAN Spark).

### Construction of the *Pseudomonas* p25.A12 tail fiber assembly gene deletion

The p25.A12△TFA mutant strain was cloned using the Gateway Cloning system with the donor vector pDONR1K18ms (Addgene plasmid #72644) and the destination vector pDEST2T18ms (Addgene plasmid #72647).

### Construction of the *Pseudomonas* p25.A12 tail fiber assembly gene overexpression rescue vector

The HTF gene was cloned into pMCSG11 (*77*) by restriction cloning (*78*). This rescue vector was transformed into p25.A12△TFA, tailocins were induced and partially purified, and killing assays were performed to assess rescue lethality.

### *In vivo* competition assays

Strain p25.C2 cultures tagged with a luciferase cassette were grown together with p25.A12 wild type or p25.A12△TFA strain. Five different ratios were used for mixing the two strains (10:10, 5:10, 3:10, 2:10, 1:10) keeping the p25.C2 strain constant at a final OD_600_ of 0.005. Bacteria were grown overnight and were diluted 1:10 the day of the experiment. Cells were grown for another 3 hours with shaking, and finally collected and mixed accordingly. The co-inoculated cells were grown in a TECAN flat white plate (TECAN, #30122300) in a final volume of 200 uL. The plates were placed in the TECAN plate reader (TECAN Spark) for 21 hours at 28°C for 55 kinetic cycles. Each cycle consisted of 15 minutes or orbital shaking at 220 rpm, resting for 5 minutes and then the luciferase-tagged (p25.C2 cells) of each well was measured. For the analysis the three highest values from each sample were used and the average luciferase value was calculated for each condition.

### *In planta* co-infections

Columbia-0 wild type *A. thaliana* plants were grown in 24-well plates (Greiner Bio-One, #6621665) under long day conditions (16 hours light, 8 hours dark) in an AR41L3 percival with 60% intensity of the SciWhite LED lights. Thirteen day-old seedlings were used for the infections. Plants were infected with bacterial suspension of the *Pseudomonas* strain p25.C2 (tagged with luciferase), and either the p25.A12 wild type or the p25.A12△TFA strain. The same five ratios as the *in vivo* assays were used, keeping the p25.C2 constant at a final OD_600_ of 0.005.

Bacteria were grown overnight and diluted 1:10 the day of the infection. Cells were grown for another three hours and finally collected and resuspended in 10mM MgSO_4_. Cells were mixed according to the ratios described above, and the seedlings were submerged in 10 mL of cell suspension for 10 minutes in a randomized manner. 850 μL were removed and the plants were placed back in the percival. Three days post infection plants were collected in 2 mL deep-well plates (Thermo Fisher^TM^ Nunc^TM^, #12-565-605) containing 1 mL MgSO_4_. Plants were ground using the Qiagen tissue lyser II (QIAGEN) and 200μL of the suspension were used to measure luciferase in the TECAN using flat white plates (TECAN, #30122300). Plants infected with MgSO_4_ or p25.C2 alone were used as controls. The experiment was repeated 3 times with similar results.

### Constructing the BarSeq Mutant Fitness Library

Methods were adapted from Wetmore et al. (*36*). Briefly, we created the p25.C2 transposon mutant library by conjugating p25.C2 with the *E. coli* conjugation donor (WM3064) harboring the pHLL250mariner transposon vector library (AMD290)(*79*). Equal cell numbers of mid-log phase p25.C2 and AMD290 were conjugated for 6 hours on 0.45 μm nitrocellulose filters (Millipore) overlaid on LB agar plates containing Diaminopimelic acid. The resuspended cells were plated on LB plates with 50 g/mL kanamycin to select for mutants. After 2 days, the kanamycin-resistant colonies were scraped into LB, the OD_600_ of the mixture was measured, and the mutant library was diluted to a starting OD_600_ of 0.2 in 250 ml of LB with 50 g/ml kanamycin. The diluted mutant library was grown at 28°C to a final OD_600_ of 1.0, added glycerol to a final volume of 10%, and stored at -80°C freezer stocks. Cells were collected for genomic DNA extraction.

### Competitive Mutant Fitness Assays

Assays were adapted from (*23*). Briefly, assays were performed in glass tubes. Partially purified tailocins were used as stressors at final concentrations of 0.05x of the stock preparation. P-buffer was used as a control. LB + Km was supplied for the mutant library as the growth medium. The tubes were incubated with shaking and mutants were harvested when the OD600 reached mid-log phase. Cells were pelleted by centrifugation (8000g, 5 min) and stored at -20°C awaiting genomic DNA extraction. Each condition was assayed in ten individual replicates.

### BarSeq and Analysis of BarSeq data

Genomic DNA was extracted and barcode PCR was performed as described in (*36*). DNA extractions were quantified with NanoDrop 1000 (Thermo Fisher). Barcode sequence data were obtained by multiplexing on a lane of HiSeq at the University of Utah High-Throughput Genomics core. Fitness data were calculated and analyzed from these reads with DESeq2 (*80*) R package and scripts can be found on our GitHub page.

### LPS isolation

The cell culture was washed two times with PBS and inactivated using a solution of 1% phenol in PBS. The cell material was suspended in water and pre-incubated to 68 °C with gentle stirring. The extraction was carried out by using the corresponding volume of 90% (vol/vol) liquefied phenol following the standard procedure (*81*) and extracted for 20 min, at 68 °C. The sample was cooled on an ice bath and centrifuged for 20 min at 4 °C, 5000 x g. The aqueous upper phase was collected, and the remaining phenol phase and cell debris were extracted two more times with water, following the same protocol. The combined aqueous phases and the final phenol phase were dialyzed against several exchanges of water, for 4 days, (MWCO).

The freeze-dried dialysates were re-suspended in a sterile buffer (10 mg/mL - 50 mM MgCl2•6H2O and 20 mM NaOAc•3H2O) and the nucleic acids were digested with Benzonase (2 µl, 18 h at 37 °C) with gentle agitation. Proteins were and digested with proteinase K (50 µg/mL - per mL of the LPS solution) by overnight digestion at 37 °C. The buffer, nucleotides, and peptides were removed by dialysis against water (12-14 kDa MWCO) at 4 °C, followed by ultracentrifugation at 100,000 x g for 16 h, at 4 °C. The LPS pellets were suspended in water, freeze-dried, and used in this work.

### DOC-PAGE of LPS samples

The purified LPS samples were resolved in PAGE (4% stacking gel and 18% resolving gel) in the presence of the deoxycholic acid buffer ((*82*)). The PAGE was visualized with a silver stain reagent kit (Bio-Rad).

### Glycosyl composition analysis by TMS method

The analysis was performed by combined gas chromatography-mass spectrometry (GC-MS) of the O-trimethylsilyl (TMS) methyl glycoside derivatives produced from the sample by acidic methanolysis followed by trimethylsilyation(*83*). GC-MS analysis of the TMS methyl glycosides was performed on an Agilent AT 7890A GC system interfaced to a 5975B MSD, using an EC-1 fused silica capillary column (30 m ’ 0.25 mm ID), and the temperature gradient was was 80°C for 2 min, then ramped to 140°C at 20°C/min with 2 min hold, and to 200°C at 2°C/min, followed by an increase to 250°C at 30°C/min with 5 min hold.

---

**Figure S1.**
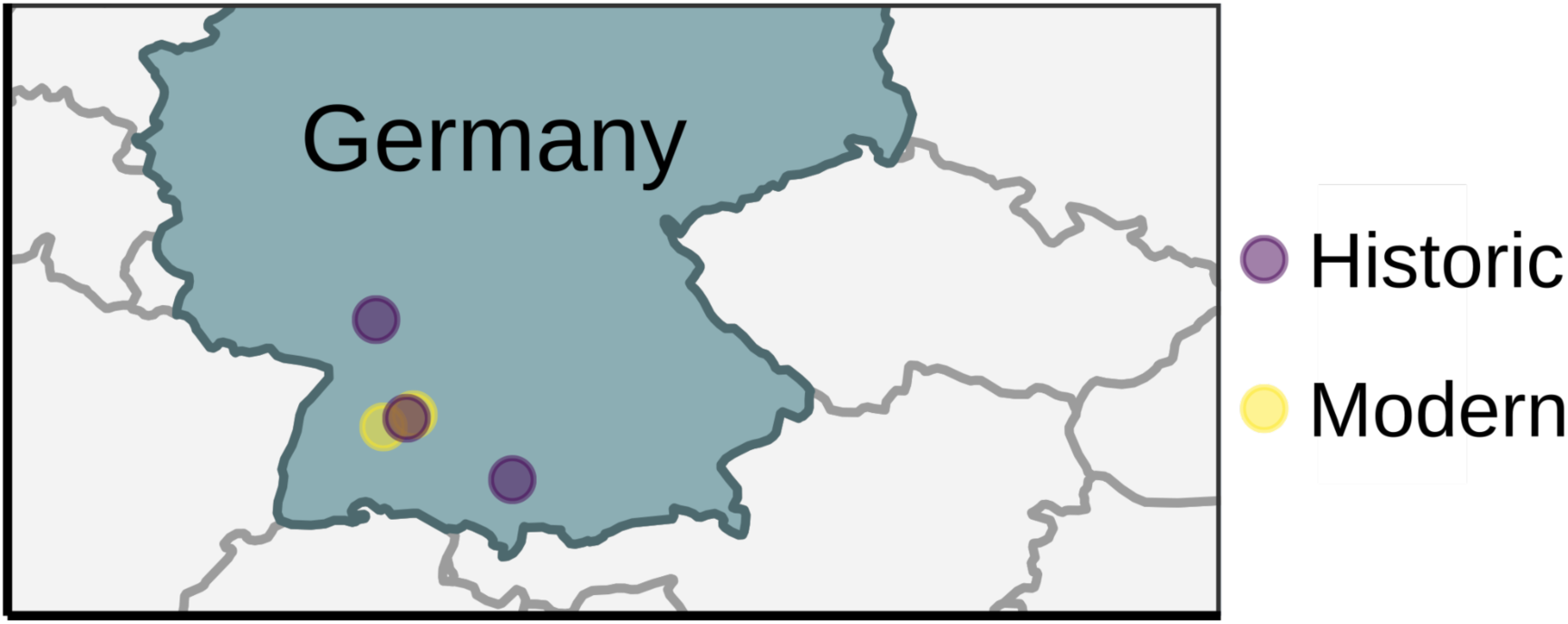
The two modern and three historic collection sites in Southwestern Germany.

**Figure S2.**
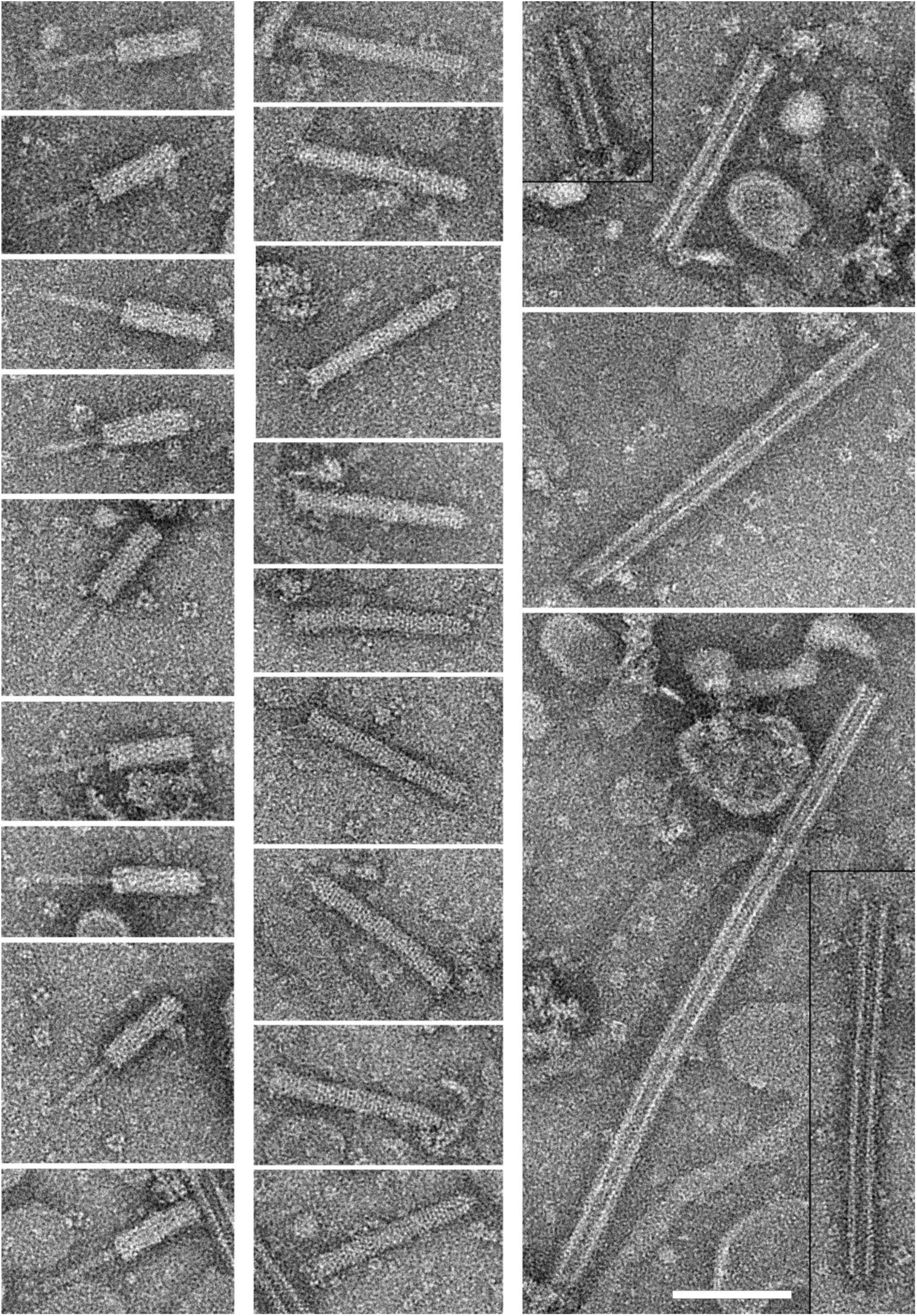
TEM images of p25.A12 contracted tailocins (left column), uncontracted tailocins (center column), and phage-like hollow tubes (right column). In the central column, the baseplate end of the tailocin faces left. The hollow tubes had highly variable lengths, average 404 (±153) nm (n = 14). Scale bar, 100 nm.

**Figure S3.**
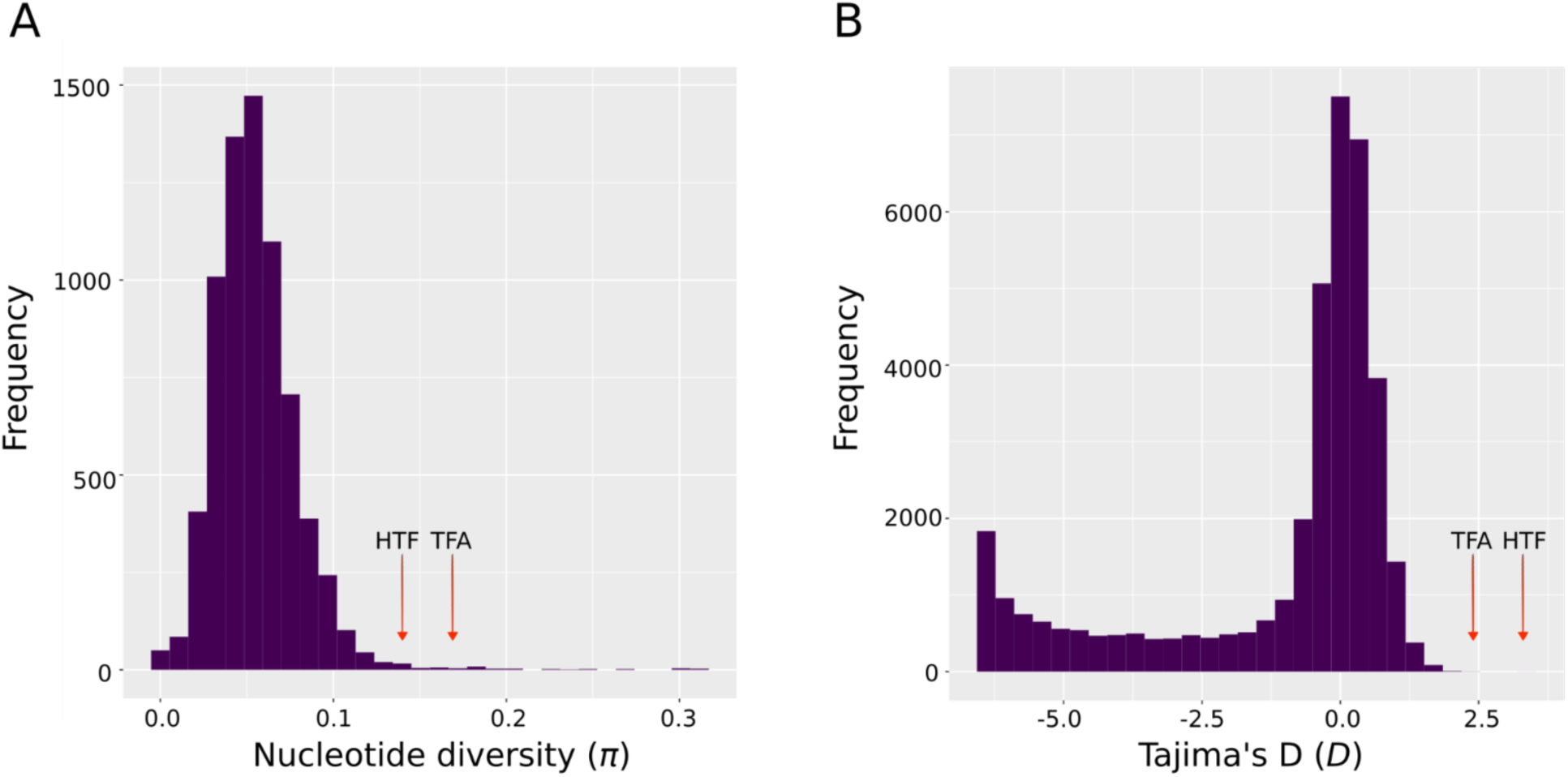
The tail fiber assembly gene and hypothetical tail fiber are evolving quicker than the rest of the bacterial genome. The distribution of **A.** nucleotide diversity (π) and **B.** Tajima’s D (D) for all core genes in the 1,399 ATUE5 genomes. The TFA and HTF genes (red arrows) have significantly higher π and D compared to the rest of the core genes.

**Figure S4.**
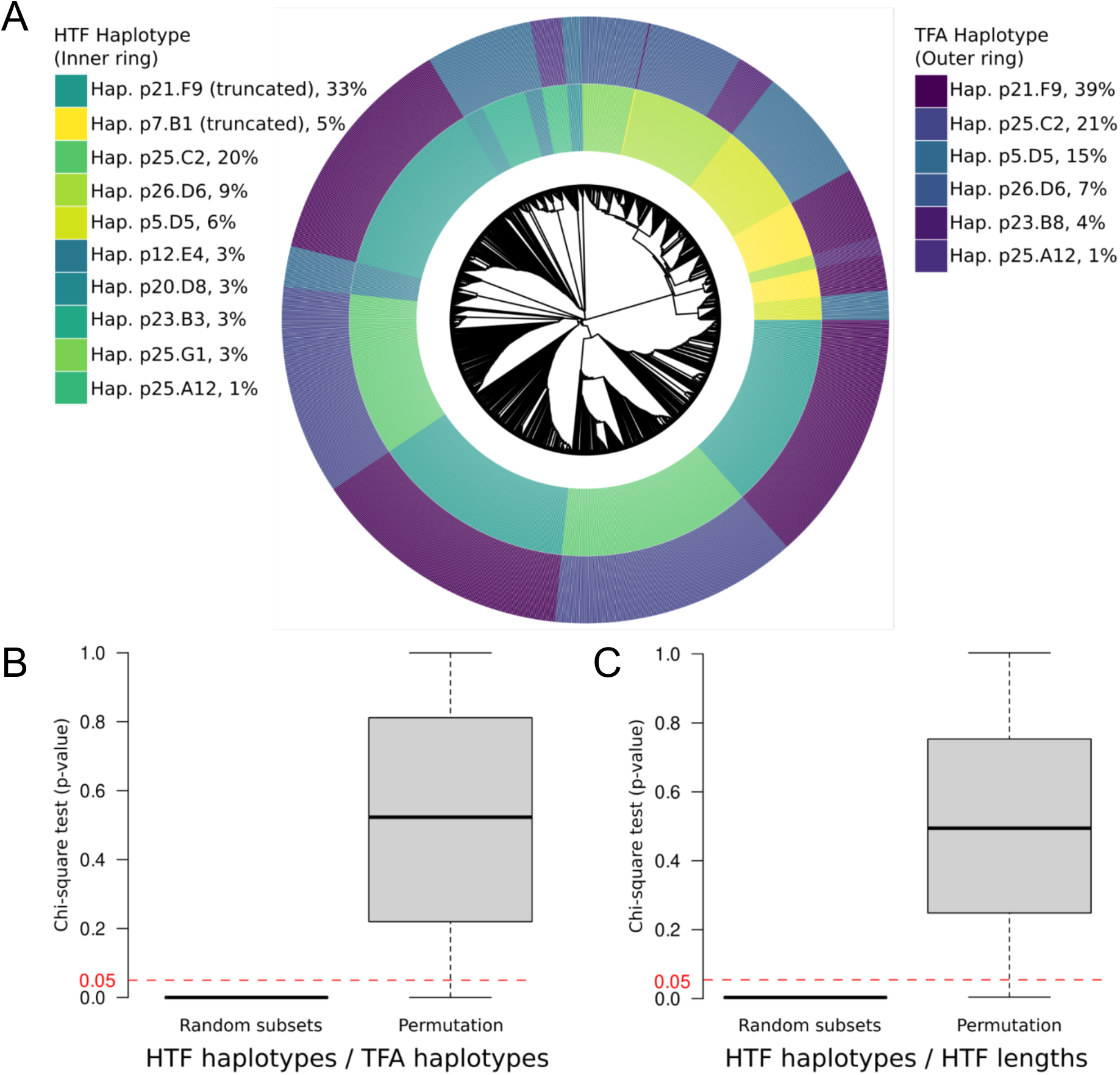
Associations between Tail Fiber Assembly (TFA) and Hypothetical Tail Fiber (HTF) haplotypes are significant. **A.** Maximum likelihood concatenated core genome phylogeny of 1,399 pathogenic pseudomonads that co-occur in the *Arabidopsis thaliana* phyllosphere with the HTF and TFA haplotypes plotted in the inner and outer rings. B-C. Each box plot represents a distribution of 1000 Chi-square test p-values drawn from random draws of pairs between TFA and HTF haplotypes and data sets for which HTF values were randomly permuted. Associations were tested between the two protein haplotypes (A) and between HTF haplotypes and TFA gene lengths (B).

**Figure S5.**
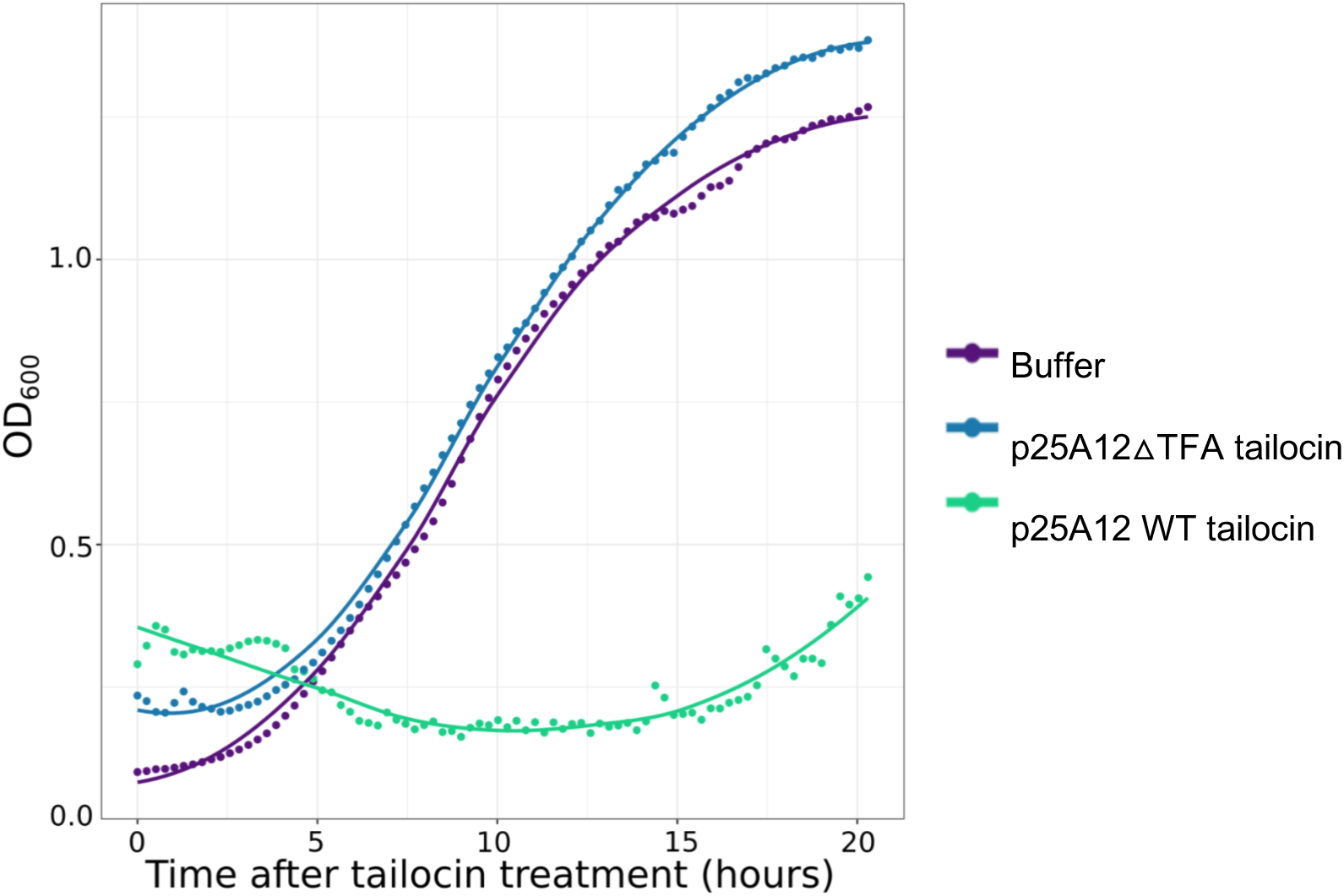
The tail fiber assembly gene was successfully knocked out and the knockout strain loses killing ability. A known sensitive strain to the tailocin (p25.C2) was grown in a Tecan for 20 hours. Treatments included p25.A12 WT purified tailocin, p25.A12 mutant purified tailocin, and a buffer control. The mean OD_600_ of two technical replicates were measured as a function of time.

**Figure S6.**
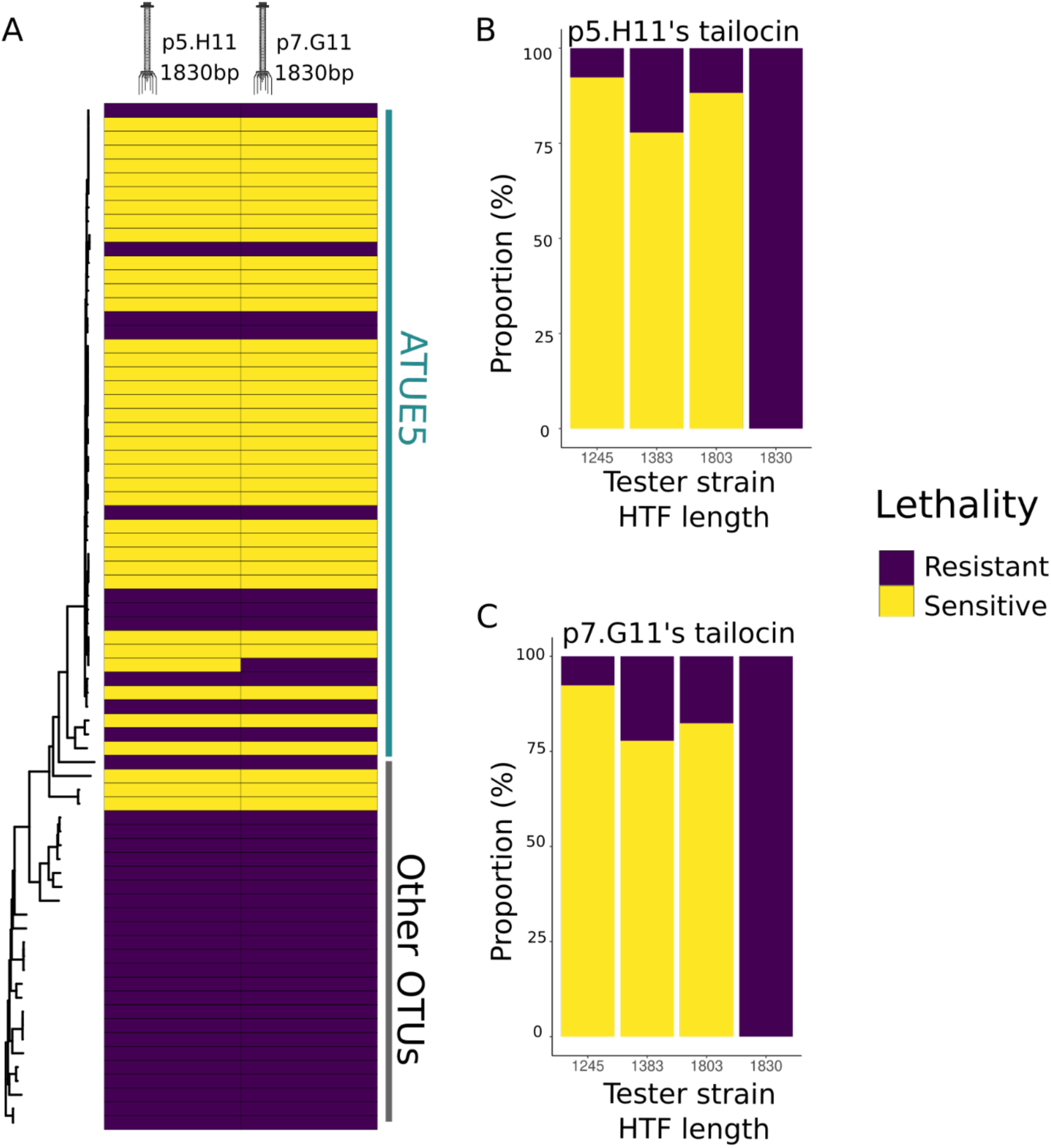
**A.** Tailocins are preferentially used for intra-lineage killing. Soft agar cultures of the *Pseudomonas* strains (rows) were challenged with viral particles extracted from cultures of two strains, p5.H11 and p7.G11 (from the 1830bp hypothetical tail fiber length haplotype, columns), in 3 technical replicates and 1 biological replicate. The phylogeny includes 83 *Pseudomonas* representative strains and are displayed according to their phylogenetic placement. Vertical lines indicate strains that belong to ATUE5 (colored in green) and other OTUs (colored in black). **B-C**. The proportion of tester strains sensitive or resistant to p5.H11’s tailocin (B) and p7.G11’s tailocin (C). Lethality significantly correlated with the tester strain’s hypothetical tail fiber length haplotype (Fisher’s Exact test, p = 10^-4^ (B), p = 10^-3^ (C).

**Figure S7.**
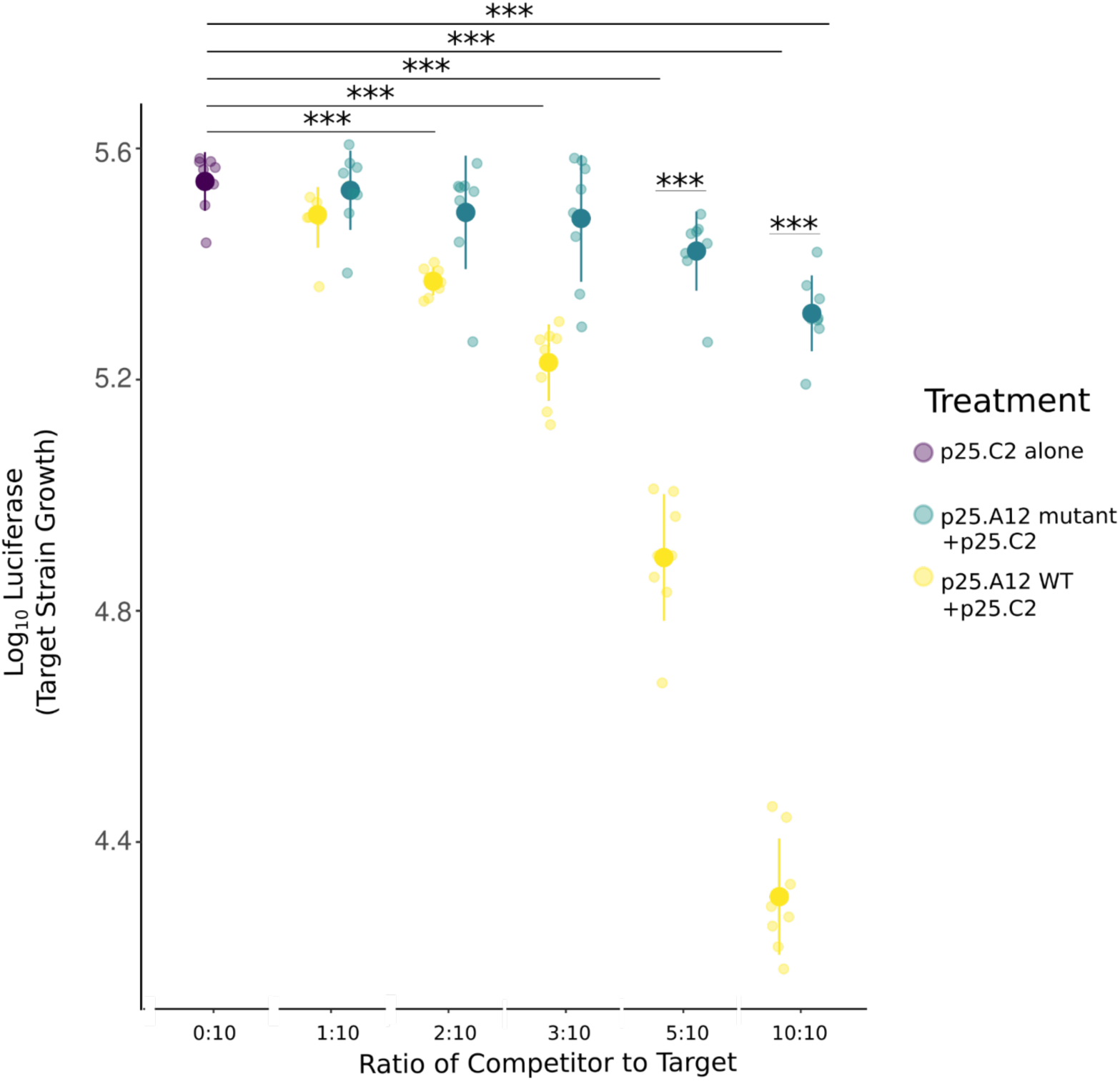
*In vivo* coinfections of p25.A12 (competitor) and p25.C2 (target, known to be sensitive to p25.A12). p25.A12 mutant competed with p25.C2 as well as p25.C2 grown alone were used as controls. Different ratios of competitor and constant amounts of target strain were used. p25.C2 grown alone as the control. ANOVA test, P-values are shown as p < 0.001 ***, p < 0.01 **, or p < 0.05 *.

**Figure S8.**
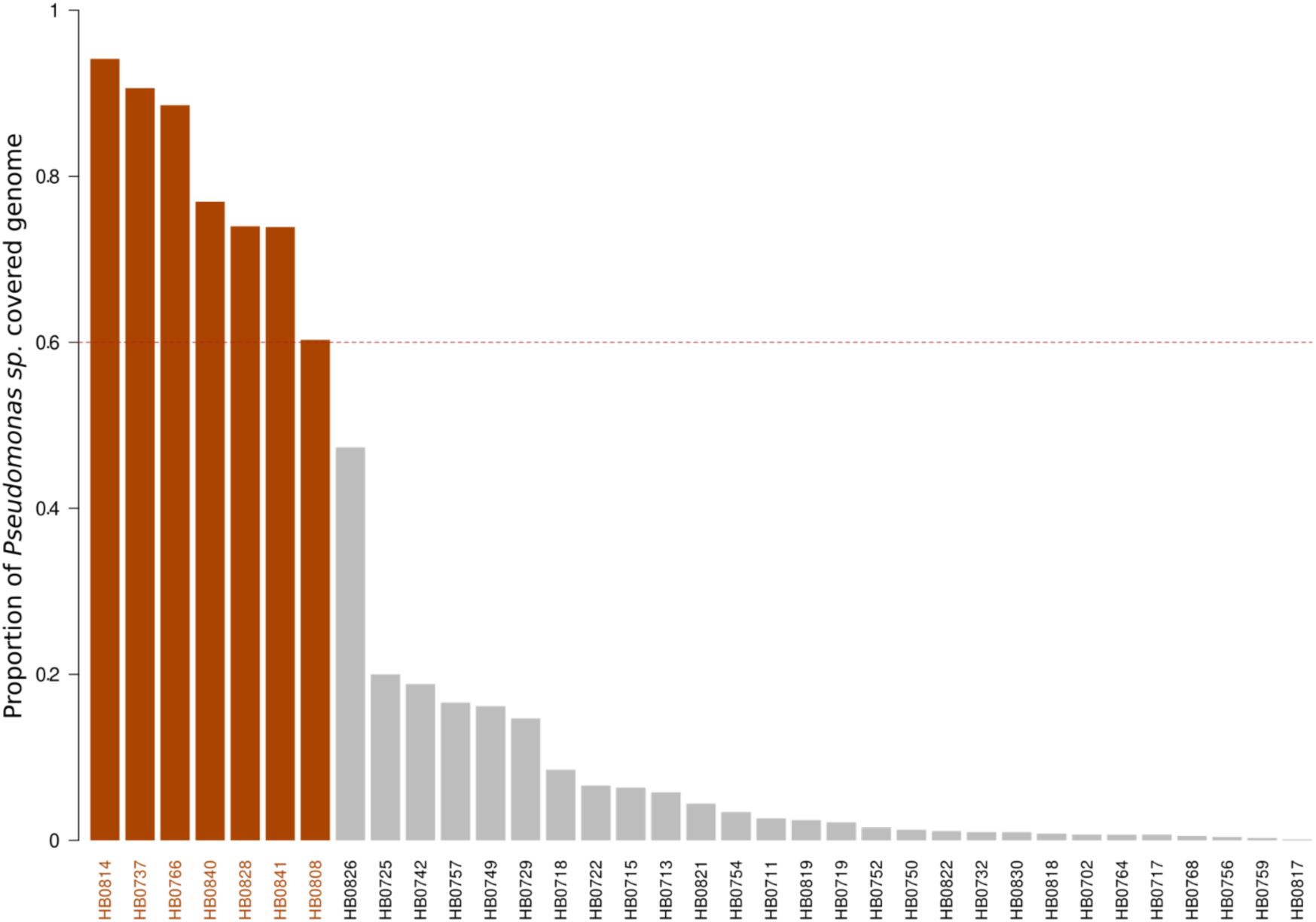
Seven historical samples cover at least 60% of the *Pseudomonas* genome. The height of the bars represent the proportion of the *Pseudomonas* sp. reference genome covered with at least one read after Mapping Quality filter value of 30. The horizontal dotted line depicts a threshold of 0.6.

**Figure S9.**
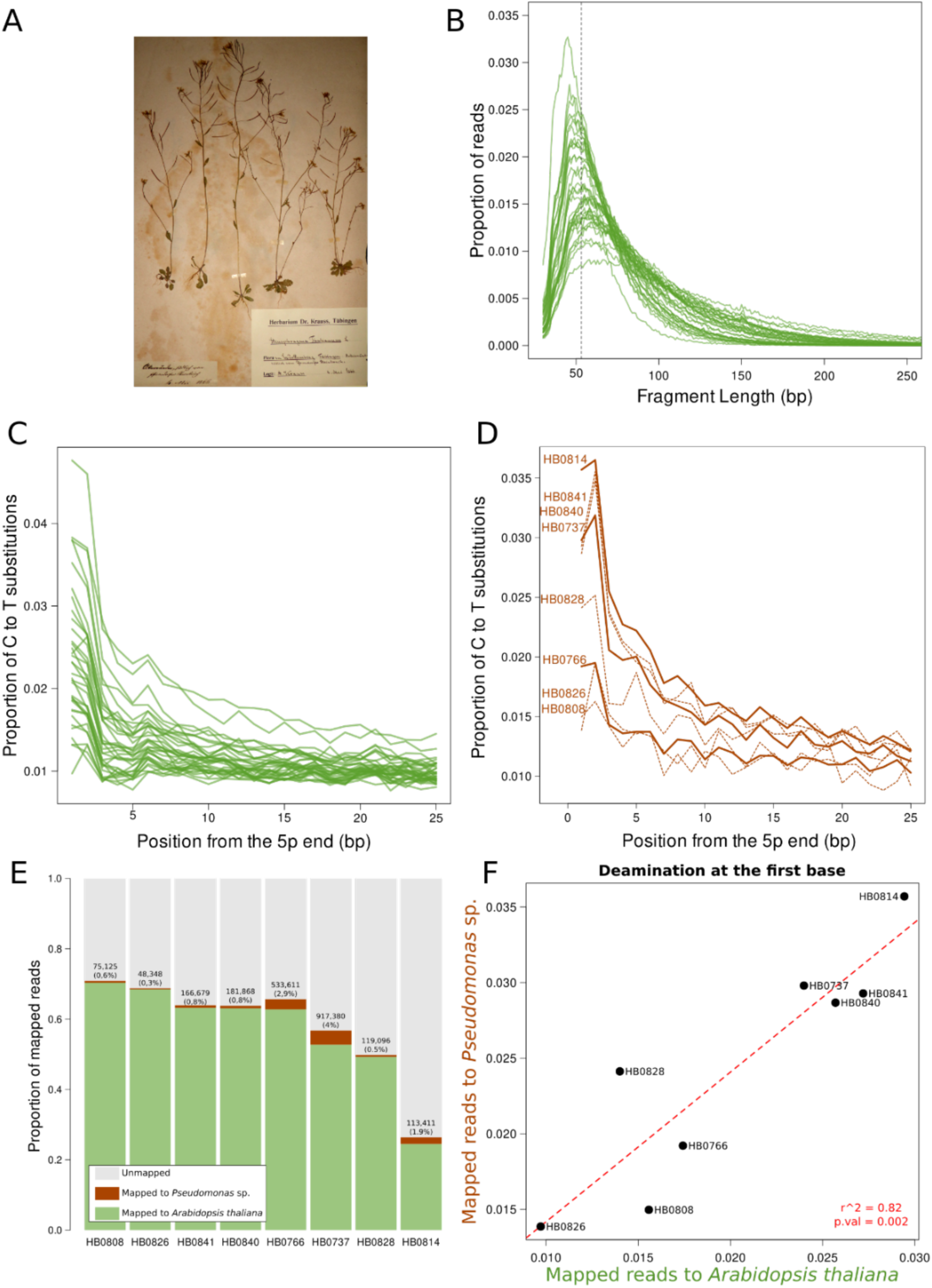
Herbarium-derived reads assigned to the host plant *Arabidopsis thaliana* and to *Pseudomonas* sp. show degradation patterns typical of ancient DNA. A. The photo shows of a representative *A. thaliana* herbarium specimen (sample HB0729) sampled in the Tübingen herbarium. **B.** Length distribution of merged sequenced reads of 35 historical herbarium samples after mapping to the *A. thaliana* reference genome. **C.** Cytosine-to-Thymine (C-to-T) substitutions at the 5’ end of reads mapped to the *A. thaliana* reference genome. **D.** C-to-T substitutions at the 5’ end of reads mapped to the *Pseudomonas viridiflava* reference genome. Only samples that covered more than 60% of the *P. viridiflava* genome were included in the analysis. The thick lines indicate samples included in the phylogenetic analyses. **E.** The bars indicate the proportion of reads assigned to *A. thaliana* or to *Pseudomonas* sp. **F.** The scatter plot displays the relationship between the proportion of C-to-T substitutions at the first base of the 5’ end between reads assigned to *A. thaliana* or to *Pseudomonas* sp. The red dotted line represents a linear model.

**Figure S10.**
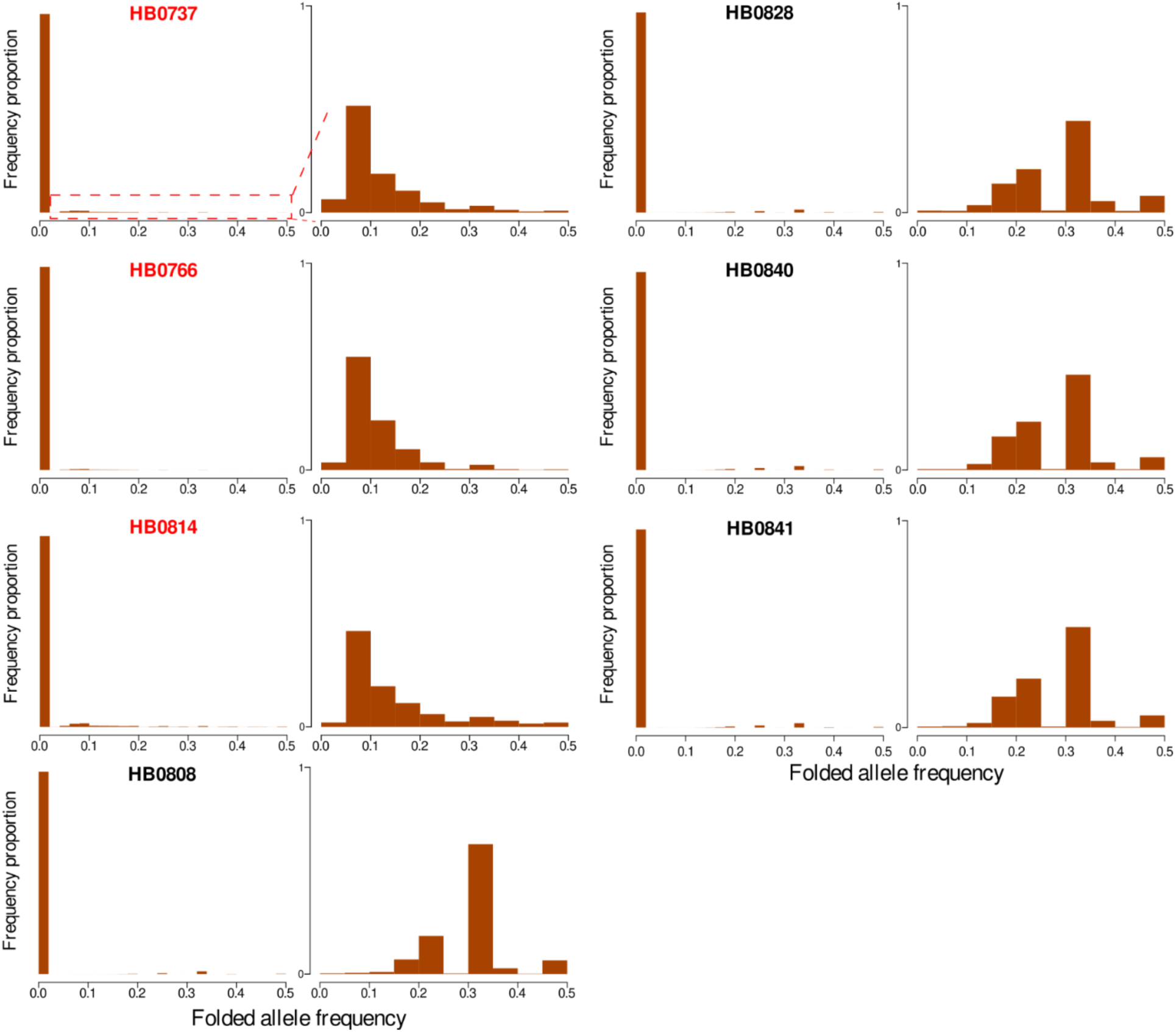
Historical *Arabidopsis thaliana* herbarium specimens are dominated by a single *Pseudomonas* sp. strain. The plots show the folded homozygosity support for each position along the *Pseudomonas viridiflava* genome with a coverage greater than 1x. A value of zero indicates that all reads support the same nucleotide (total homozygosity), where a value of 0.5 indicates maximum heterozygosity with only half of the reads supporting a particular nucleotide. Each row shows the folded homozygosity support for each sample. The left column shows all bins, whereas the bar indicating total homozygosity was removed in the right column.

**Figure S11.**
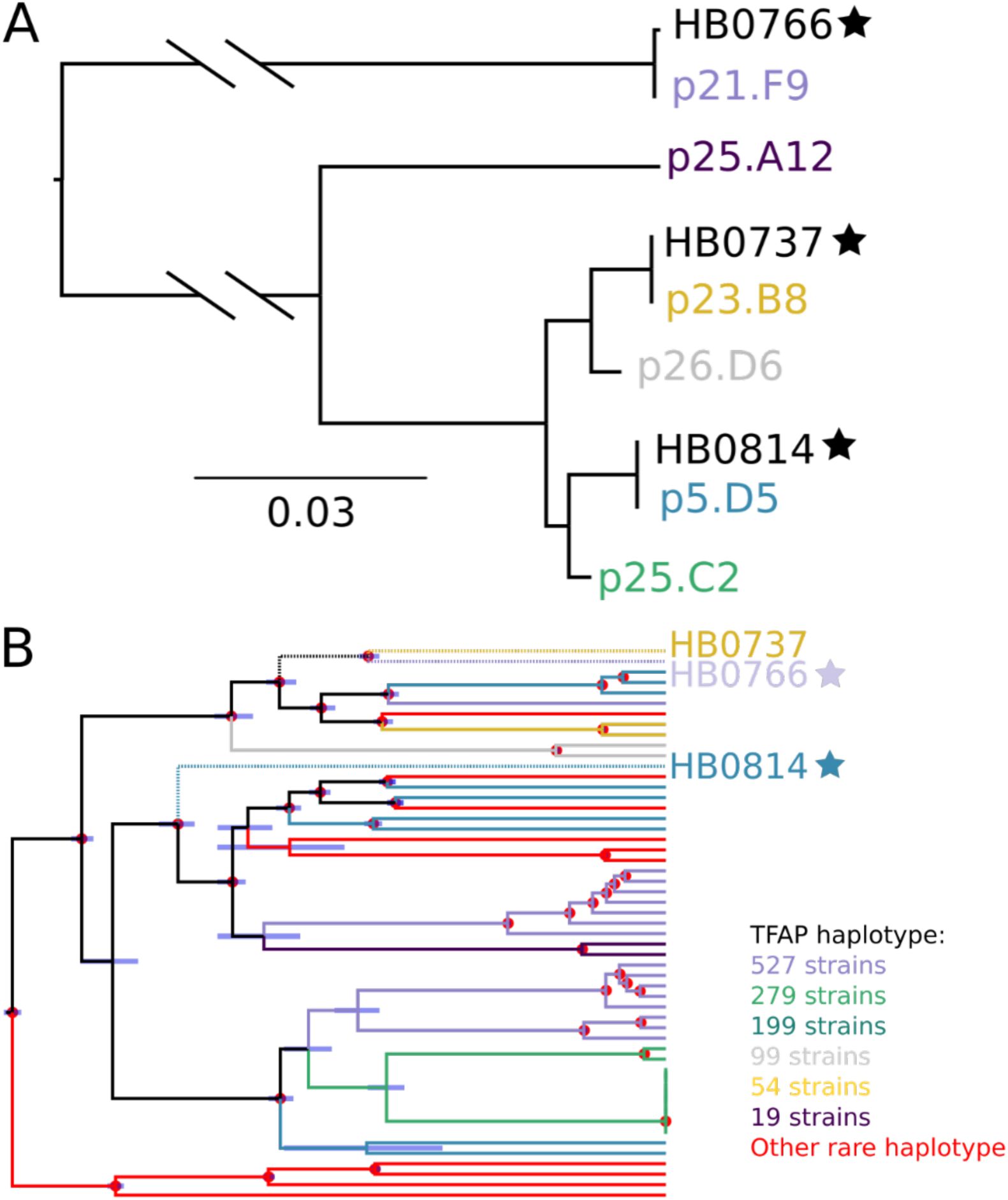
**A.** Neighbor-Joining tree of the tail fiber assembly gene translated sequence. Historical samples (HB) are placed in the context of the most common haplotypes. **B.** Bayesian tip-date calibrated phylogeny representing the evolutionary relationship between historical and modern *Pseudomonas* sp. strains. The tips and branches are colored based on the tail fiber assembly gene haplotype. Historical samples are shown with stars. The node bars represent the 95% Highest Posterior Density Intervals of the estimated time and the nodes marked with red dots represent those with posterior probability of 1.

**Table S1.**
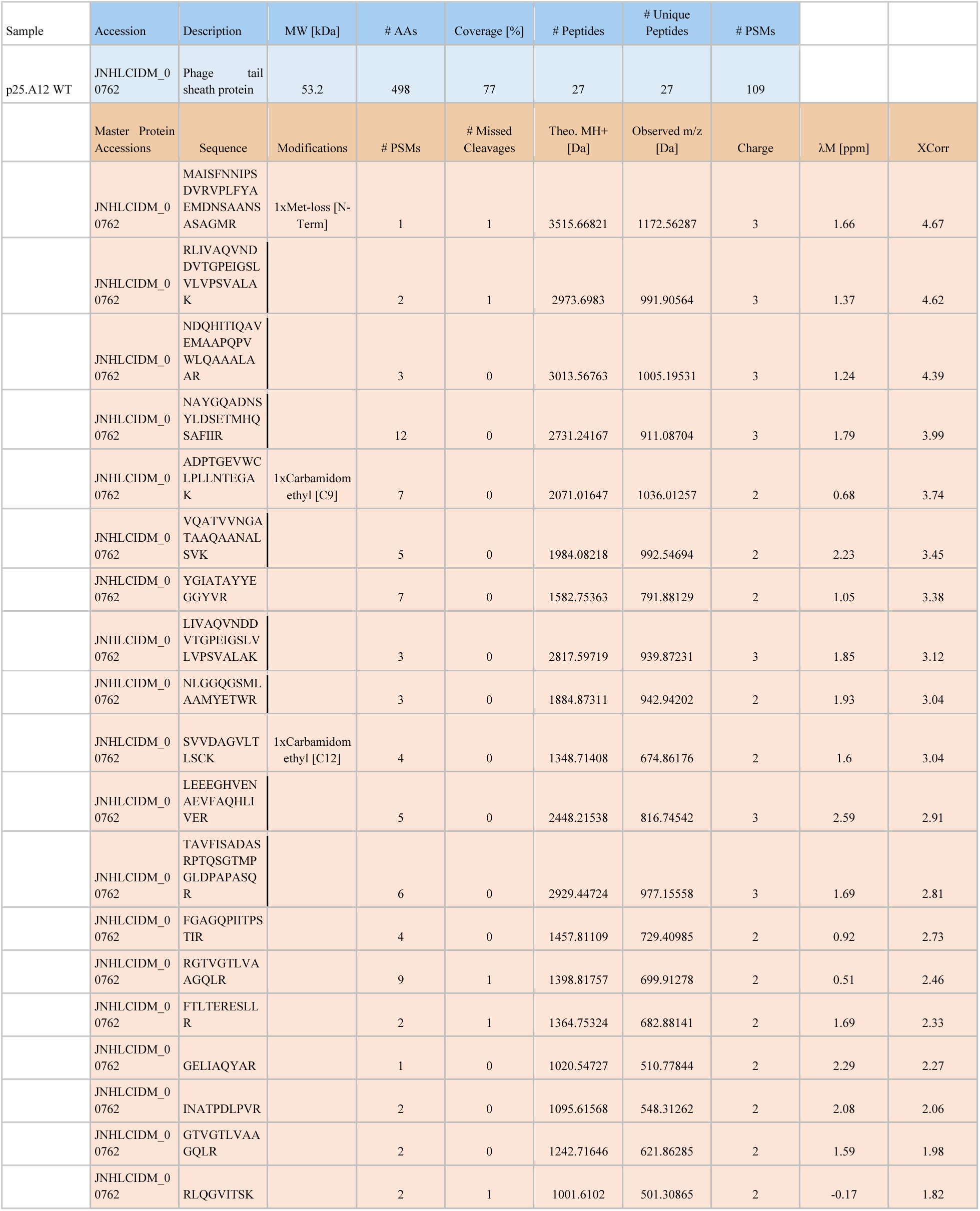
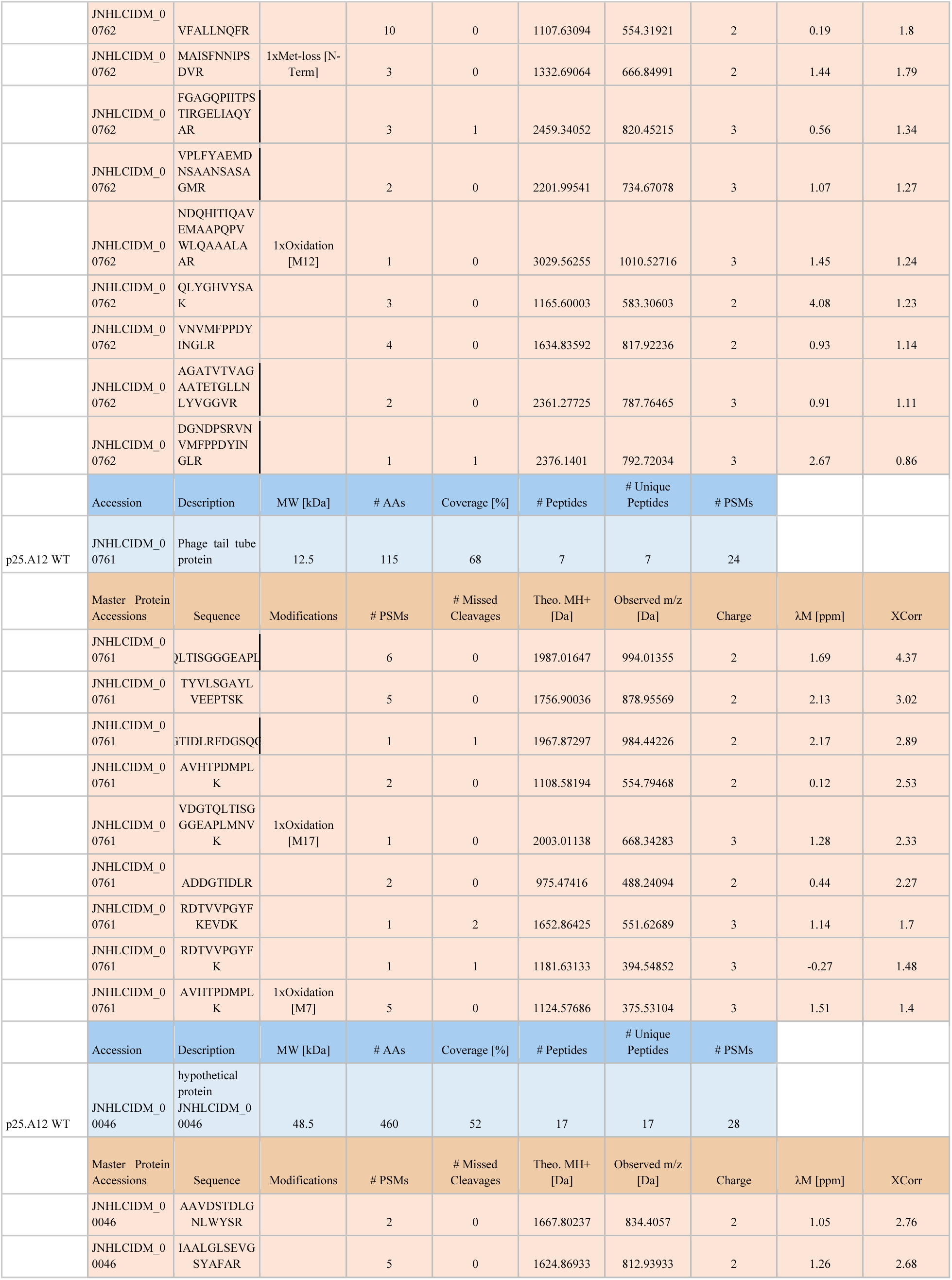
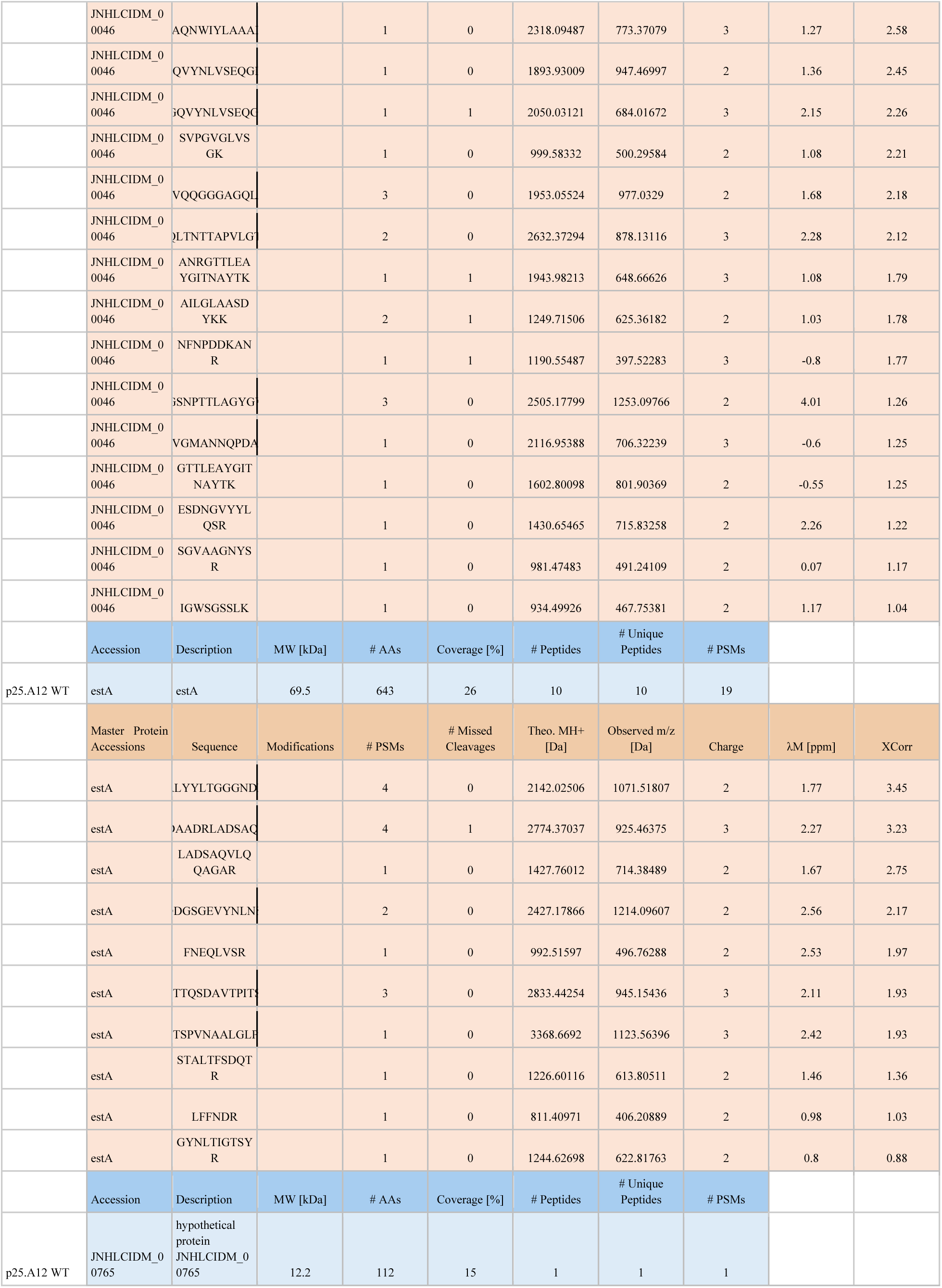
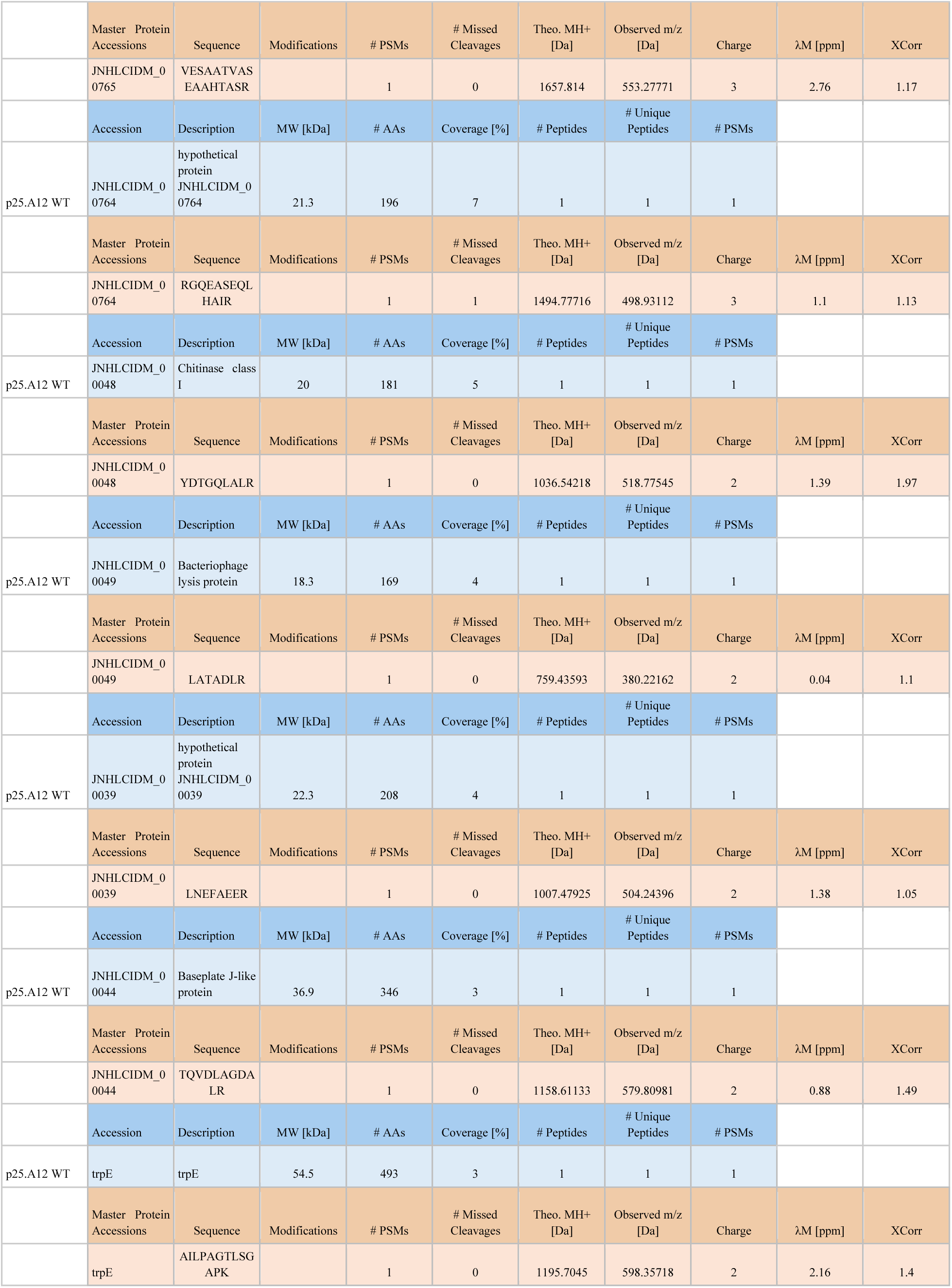
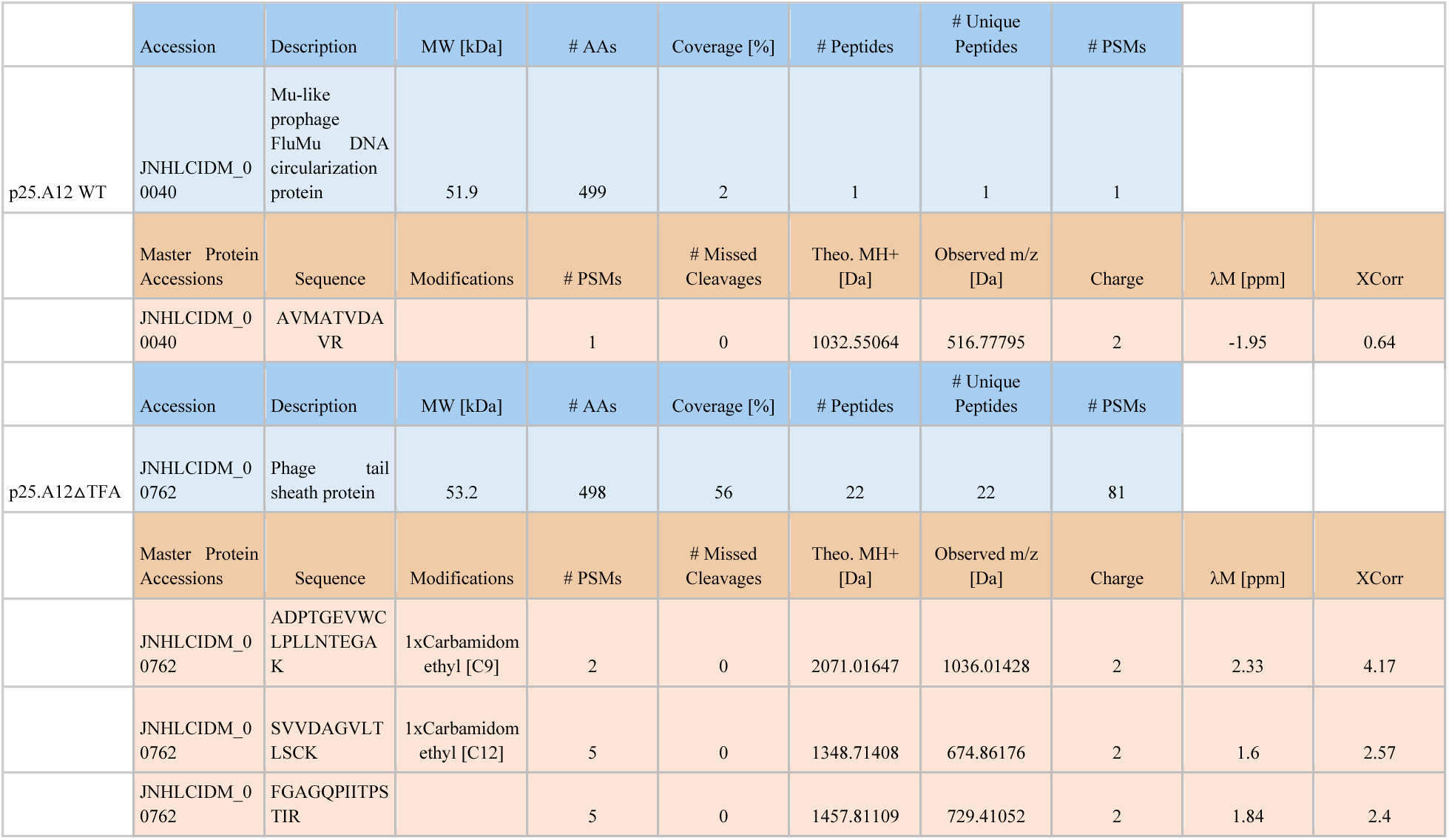
Untargeted data dependent LC/MS/MS Proteomics data for p25.A12 WT and p25.A12△TFA tailocin partially purified extracts.

**Table S2:** Relative mole percentage of monosaccharide residues detected by GC/MS.

| Residue/Strain | Mole % |  |  |  |
| --- | --- | --- | --- | --- |
|  | p21.F1 | p21.F9 | p25.A12 | p25.C2 |
| **Rha** | **63.4** | **11.1** | **9.6** | **72.3** |
| **Man** | **0.1** | **0.2** | **0.1** | **0.1** |
| **Gal** | **0.0** | **0.0** | **0.1** | **0.0** |
| **Glc** | **21.7** | **56.5** | **75.4** | **14.5** |
| **Hep** | **0.2** | **Tr.** | **0.0** | **0.0** |
| **Kdo** | **3.0** | **11.4** | **1.2** | **0.3** |
| **FucNAc** | **5.8** | **0.0** | **0.0** | **5.1** |
| **QuiNAc** | **1.1** | **0.0** | **0.6** | **0.8** |
| **ManNAc** | **0.3** | **1.9** | **0.9** | **0.6** |
| **GalNAc** | **0.3** | **1.1** | **0.7** | **0.3** |
| **GlcNAc** | **4.2** | **17.6** | **11.6** | **6.0** |
| **% CHO** | **60.6%** | **22.5%** | **31%** | **52%** |

Tr.- trace

**Table S3:** Historic and modern sample metadata.

\* Type: (H)istoric ; (M)odern
| ACCESSION_ID | ENA_ID | TYPE | COLLECTION_YEAR | COUNTRY | LOCATION | LAT | LON | Pseudomonas_Genome_Coverage_Prop. | Pseudomonas_Genome_Average_Depth_MQ30 | Used_for_authentication_analyses | downstream_analysis |
| --- | --- | --- | --- | --- | --- | --- | --- | --- | --- | --- | --- |
| HB0702 | SRR21814520 | H | 1905 | DEU | Mellendorf_near_Buttlar | 52.5470675 | 9.7299326 | 0.00701769 | 0.00629547 | NO | NO |
| HB0711 | SRR21814519 | H | 1840 | DEU | NA | NA | NA | 0.0266046 | 0.0199101 | NO | NO |
| HB0713 | SRR21814508 | H | 1826 | DEU | Esslingen | 48.7427584 | 9.3071685 | 0.057587 | 0.0515586 | NO | NO |
| HB0715 | SRR21814497 | H | 1828 | DEU | Esslingen | 48.7427584 | 9.3071685 | 0.0634669 | 0.0581094 | NO | NO |
| HB0717 | SRR21814491 | H | 1826 | DEU | Alpirsbach | 48.3456021 | 8.4033079 | 0.00664118 | 0.00581024 | NO | NO |
| HB0718 | SRR21814490 | H | 1890 | DEU | Tuebingen | 48.5236164 | 9.0535531 | 0.0850372 | 0.0709387 | NO | NO |
| HB0719 | SRR21814489 | H | 1890 | DEU | Tuebingen | 48.5236164 | 9.0535531 | 0.0216538 | 0.0139758 | NO | NO |
| HB0722 | SRR21814488 | H | 1891 | DEU | Altburg | 48.6619431 | 8.6926366 | 0.0659517 | 0.0522472 | NO | NO |
| HB0725 | SRR21814487 | H | 1888 | DEU | Tuebingen | 48.5236164 | 9.0535531 | 0.199974 | 0.197268 | NO | NO |
| HB0729 | SRR21814486 | H | 1866 | DEU | Tuebingen | 48.5236164 | 9.0535531 | 0.146878 | 0.136151 | NO | NO |
| HB0732 | SRR21814518 | H | 1932 | DEU | Rastatt | 49.0532906 | 8.5245089 | 0.0100316 | 0.0070049 | NO | NO |
| HB0737 | SRR21814517 | H | 1937 | DEU | Tuebingen | 48.5236164 | 9.0535531 | 0.906105 | 8.75871 | YES | YES |
| HB0742 | SRR21814516 | H | 1953 | DEU | Lichtenstein | 48.41996913 | 9.26630222 | 0.187988 | 0.162055 | NO | NO |
| HB0749 | SRR21814515 | H | 1849 | DEU | near_Hohenheim | 48.7118 | 9.2113 | 0.161478 | 0.156122 | NO | NO |
| HB0750 | SRR21814514 | H | 1957 | DEU | Boeblingen | 48.684969 | 9.0113444 | 0.012546 | 0.010024 | NO | NO |
| HB0752 | SRR21814513 | H | 1952 | DEU | Eberspiel_in_the_district_of_Calw | 48.7415506 | 8.6842699 | 0.0150715 | 0.0118002 | NO | NO |
| HB0754 | SRR21814512 | H | 1951 | DEU | District_of_Calw | 48.7153344 | 8.7381796 | 0.033677 | 0.028471 | NO | NO |
| HB0756 | SRR21814511 | H | 1951 | DEU | Wendlingen | 50.8021737 | 7.7130698 | 0.00412007 | 0.00351381 | NO | NO |
| HB0757 | SRR21814510 | H | 1951 | DEU | Wendlingen | 50.8021737 | 7.7130698 | 0.165843 | 0.100545 | NO | NO |
| HB0759 | SRR21814509 | H | 1893 | DEU | Wohlmuthausen | 50.5727241 | 10.21353915 | 0.00280539 | 0.00201518 | NO | NO |
| HB0764 | SRR21814507 | H | 1905 | DEU | Mengen | 48.049796 | 9.3316096 | 0.00683053 | 0.00540983 | NO | NO |
| HB0766 | SRR21814506 | H | 1846 | DEU | Monastery_for<br>est_Hohenzoll<br>ern | 47.9724751 | 10.3005044 | 0.885642 | 7.63767 | YES | YES |
| HB0768 | SRR21814505 | H | 1894 | DEU | Todtnau | 47.8303378 | 7.9452378 | 0.00528544 | 0.00427861 | NO | NO |
| HB0808 | SRR21814504 | H | 1817 | DEU | Stuttgart_Moe<br>hringen | 48.7784485 | 9.1800132 | 0.60285 | 0.974844 | YES | NO |
| HB0814 | SRR21814503 | H | 1882 | DEU | Heidelberg | 49.4093582 | 8.694724 | 0.941364 | 8.13508 | YES | YES |
| HB0817 | SRR21814502 | H | 1934 | DEU | Gundelwangen<br>_near_Bonndo<br>rf | 47.8426163 | 8.2864051 | 0.00113424 | 0.00213973 | NO | NO |
| HB0818 | SRR21814501 | H | 1928 | DEU | Langenau_nea<br>r_Ulm | 48.7276553 | 8.0150864 | 0.00812181 | 0.00744537 | NO | NO |
| HB0819 | SRR21814500 | H | 1928 | DEU | Langenau_nea<br>r_Ulm | 48.7276553 | 8.0150864 | 0.0241737 | 0.0120816 | NO | NO |
| HB0821 | SRR21814499 | H | 1874 | DEU | Winnenden | 48.8754571 | 9.3978478 | 0.0443252 | 0.0396616 | NO | NO |
| HB0822 | SRR21814498 | H | 1936 | DEU | Hohentwiel | 47.7617515 | 8.8348709 | 0.0109147 | 0.0104388 | NO | NO |
| HB0826 | SRR21814496 | H | 1948 | DEU | Abtsgmuend | 48.8936471 | 10.0027636 | 0.473287 | 0.665255 | NO | NO |
| HB0828 | SRR21814495 | H | 1890 | DEU | Schramberg | 48.225478 | 8.3852168 | 0.739741 | 1.6142 | YES | NO |
| HB0830 | SRR21814494 | H | 1862 | DEU | near_Beimerst<br>etten | 48.4840643 | 9.9836241 | 0.00970948 | 0.00894266 | NO | NO |
| HB0840 | SRR21814493 | H | 1851 | DEU | Sulz_am_Nec<br>kar | 48.3617509 | 8.6314329 | 0.769394 | 1.61748 | YES | NO |
| HB0841 | SRR21814492 | H | 1875 | DEU | Freudenstadt_i<br>n_the_Black_<br>Forest | 48.4637727 | 8.4111727 | 0.738722 | 1.44545 | YES | NO |
| p12.A11 | NA | M | 2018 | DEU | Eyach | 48.446111 | 8.781611 | 0.955211 | 43.2805 | NA | YES |
| p12.A9 | NA | M | 2018 | DEU | Eyach | 48.446111 | 8.781611 | 0.385298 | 2.68517 | NA | NO |
| p12.E2 | NA | M | 2018 | DEU | Eyach | 48.446111 | 8.781611 | 0.946364 | 124.361 | NA | YES |
| p12.F2 | NA | M | 2018 | DEU | Det-2 | 48.556255 | 9.135424 | 0.941151 | 61.1632 | NA | YES |
| p12.G7 | NA | M | 2018 | DEU | Det-2 | 48.556255 | 9.135424 | 0.942483 | 26.1727 | NA | YES |
| p12.H7 | NA | M | 2018 | DEU | Det-2 | 48.556255 | 9.135424 | 0.703408 | 7.44366 | NA | NO |
| p13.C1 | NA | M | 2018 | DEU | Eyach | 48.446111 | 8.781611 | 0.919667 | 91.4711 | NA | YES |
| p13.C7 | NA | M | 2018 | DEU | Eyach | 48.446111 | 8.781611 | 0.915668 | 30.2041 | NA | NO |
| p13.D10 | NA | M | 2018 | DEU | Eyach | 48.446111 | 8.781611 | 0.917782 | 83.4618 | NA | YES |
| p13.D5 | NA | M | 2018 | DEU | Eyach | 48.446111 | 8.781611 | 0.373545 | 3.2645 | NA | NO |
| p13.F1 | NA | M | 2018 | DEU | Eyach | 48.446111 | 8.781611 | 0.548638 | 9.89508 | NA | NO |
| p13.F3 | NA | M | 2018 | DEU | Eyach | 48.446111 | 8.781611 | 0.930129 | 180.182 | NA | YES |
| p13.F5 | NA | M | 2018 | DEU | Eyach | 48.446111 | 8.781611 | 0.486993 | 7.68403 | NA | NO |
| p20.B10 | NA | M | 2018 | DEU | Eyach | 48.446111 | 8.781611 | 0.362051 | 6.0333 | NA | NO |
| p20.D4 | NA | M | 2018 | DEU | Eyach | 48.446111 | 8.781611 | 0.965996 | 26.275 | NA | YES |
| p20.F10 | NA | M | 2018 | DEU | Eyach | 48.446111 | 8.781611 | 0.951949 | 27.3486 | NA | YES |
| p20.G9 | NA | M | 2018 | DEU | Eyach | 48.446111 | 8.781611 | 0.951606 | 25.8164 | NA | YES |
| p21.A8 | NA | M | 2018 | DEU | Eyach | 48.446111 | 8.781611 | 0.951989 | 50.4423 | NA | YES |
| p21.E3 | NA | M | 2018 | DEU | Eyach | 48.446111 | 8.781611 | 0.912764 | 42.5976 | NA | YES |
| p21.F1 | NA | M | 2018 | DEU | Eyach | 48.446111 | 8.781611 | 0.921125 | 54.2176 | NA | YES |
| p21.F9 | NA | M | 2018 | DEU | Eyach | 48.446111 | 8.781611 | 0.951456 | 43.9553 | NA | YES |
| p22.A8 | NA | M | 2018 | DEU | Eyach | 48.446111 | 8.781611 | 0.910773 | 42.5346 | NA | YES |
| p22.B5 | NA | M | 2018 | DEU | Eyach | 48.446111 | 8.781611 | 0.908795 | 31.8141 | NA | YES |
| p22.C1 | NA | M | 2018 | DEU | Eyach | 48.446111 | 8.781611 | 0.910697 | 44.4703 | NA | YES |
| p22.D1 | NA | M | 2018 | DEU | Eyach | 48.446111 | 8.781611 | 0.93509 | 9.86993 | NA | YES |
| p22.D4 | NA | M | 2018 | DEU | Eyach | 48.446111 | 8.781611 | 0.901368 | 6.48617 | NA | YES |
| p23.A3 | NA | M | 2018 | DEU | Det-2 | 48.556255 | 9.135424 | 0.922147 | 90.3312 | NA | YES |
| p23.A5 | NA | M | 2018 | DEU | Det-2 | 48.556255 | 9.135424 | 0.355851 | 5.69028 | NA | NO |
| p23.B4 | NA | M | 2018 | DEU | Det-2 | 48.556255 | 9.135424 | 0.353754 | 10.3529 | NA | NO |
| p23.B8 | NA | M | 2018 | DEU | Pfrondorf2 | 48.561087 | 9.109294 | 0.915323 | 25.8754 | NA | YES |
| p24.B5 | NA | M | 2018 | DEU | Eyach | 48.446111 | 8.781611 | 0.906998 | 24.3191 | NA | YES |
| p24.H2 | NA | M | 2018 | DEU | Eyach | 48.446111 | 8.781611 | 0.919194 | 36.961 | NA | YES |
| p25.A12 | NA | M | 2018 | DEU | Eyach | 48.446111 | 8.781611 | 0.904389 | 8.27369 | NA | YES |
| p25.B2 | NA | M | 2018 | DEU | Eyach | 48.446111 | 8.781611 | 0.99527 | 9.73957 | NA | YES |
| p25.C11 | NA | M | 2018 | DEU | Eyach | 48.446111 | 8.781611 | 0.936486 | 7.67613 | NA | NO |
| p25.C2 | NA | M | 2018 | DEU | Eyach | 48.446111 | 8.781611 | 0.999957 | 38.9734 | NA | YES |
| p25.D2 | NA | M | 2018 | DEU | Eyach | 48.446111 | 8.781611 | 0.997331 | 11.0366 | NA | YES |
| p26.B7 | NA | M | 2018 | DEU | Pfrondorf2 | 48.561087 | 9.109294 | 0.918811 | 18.5831 | NA | YES |
| p26.C10 | NA | M | 2018 | DEU | Det-2 | 48.556255 | 9.135424 | 0.532614 | 7.27294 | NA | NO |
| p26.D6 | NA | M | 2018 | DEU | Eyach | 48.446111 | 8.781611 | 0.907977 | 10.1768 | NA | YES |
| p26.E7 | NA | M | 2018 | DEU | Pfrondorf2 | 48.561087 | 9.109294 | 0.915118 | 26.4647 | NA | YES |
| p26.F6 | NA | M | 2018 | DEU | Pfrondorf2 | 48.561087 | 9.109294 | 0.263831 | 1.09767 | NA | NO |
| p27.C5 | NA | M | 2018 | DEU | Pfrondorf2 | 48.561087 | 9.109294 | 0.33096 | 4.98267 | NA | NO |
| p27.D6 | NA | M | 2018 | DEU | Pfrondorf2 | 48.561087 | 9.109294 | 0.915116 | 62.9111 | NA | YES |
| p27.F2 | NA | M | 2018 | DEU | Pfrondorf2 | 48.561087 | 9.109294 | 0.908864 | 30.4347 | NA | YES |
| p3.A3 | NA | M | 2018 | DEU | Eyach | 48.446111 | 8.781611 | 0.999479 | 12.1995 | NA | YES |
| p3.F12 | NA | M | 2018 | DEU | Eyach | 48.446111 | 8.781611 | 0.999766 | 64.6266 | NA | YES |
| p3.F8 | NA | M | 2018 | DEU | Eyach | 48.446111 | 8.781611 | 0.910145 | 14.5013 | NA | NO |
| p3.G11 | NA | M | 2018 | DEU | Eyach | 48.446111 | 8.781611 | 0.405107 | 35.7165 | NA | NO |
| p3.G9 | NA | M | 2018 | DEU | Eyach | 48.446111 | 8.781611 | 0.936757 | 40.7657 | NA | YES |
| p4.A6 | NA | M | 2018 | DEU | Eyach | 48.446111 | 8.781611 | 0.999359 | 11.257 | NA | YES |
| p4.A8 | NA | M | 2018 | DEU | Eyach | 48.446111 | 8.781611 | 0.317566 | 8.57523 | NA | NO |
| p4.C5 | NA | M | 2018 | DEU | Eyach | 48.446111 | 8.781611 | 0.281958 | 2.19208 | NA | NO |
| p4.D11 | NA | M | 2018 | DEU | Eyach | 48.446111 | 8.781611 | 0.329928 | 7.72875 | NA | NO |
| p4.D2 | NA | M | 2018 | DEU | Eyach | 48.446111 | 8.781611 | 0.903089 | 43.5342 | NA | YES |
| p4.E5 | NA | M | 2018 | DEU | Eyach | 48.446111 | 8.781611 | 0.905147 | 9.07888 | NA | YES |
| p4.E6 | NA | M | 2018 | DEU | Eyach | 48.446111 | 8.781611 | 0.907851 | 26.4274 | NA | YES |
| p4.H3 | NA | M | 2018 | DEU | Det-2 | 48.556255 | 9.135424 | 0.359271 | 11.1964 | NA | NO |
| p5.A5 | NA | M | 2018 | DEU | Eyach | 48.446111 | 8.781611 | 0.569164 | 20.9644 | NA | NO |
| p5.C1 | NA | M | 2018 | DEU | Eyach | 48.446111 | 8.781611 | 0.940631 | 83.5842 | NA | YES |
| p5.C3 | NA | M | 2018 | DEU | Eyach | 48.446111 | 8.781611 | 0.911686 | 61.8474 | NA | YES |
| p5.D5 | NA | M | 2018 | DEU | Eyach | 48.446111 | 8.781611 | 0.908126 | 47.1102 | NA | YES |
| p5.F2 | NA | M | 2018 | DEU | Eyach | 48.446111 | 8.781611 | 0.355109 | 10.8067 | NA | NO |
| p5.H11 | NA | M | 2018 | DEU | Eyach | 48.446111 | 8.781611 | 0.912223 | 24.6014 | NA | YES |
| p6.A10 | NA | M | 2018 | DEU | Eyach | 48.446111 | 8.781611 | 0.967914 | 86.1443 | NA | YES |
| p6.B5 | NA | M | 2018 | DEU | Eyach | 48.446111 | 8.781611 | 0.377432 | 8.17665 | NA | NO |
| p6.B9 | NA | M | 2018 | DEU | Eyach | 48.446111 | 8.781611 | 0.915641 | 40.9034 | NA | NO |
| p6.D10 | NA | M | 2018 | DEU | Eyach | 48.446111 | 8.781611 | 0.362251 | 6.29957 | NA | NO |
| p6.E9 | NA | M | 2018 | DEU | Eyach | 48.446111 | 8.781611 | 0.348102 | 5.45088 | NA | NO |
| p6.F1 | NA | M | 2018 | DEU | Det-2 | 48.556255 | 9.135424 | 0.441901 | 14.1066 | NA | NO |
| p6.G2 | NA | M | 2018 | DEU | Det-2 | 48.556255 | 9.135424 | 0.591204 | 22.8581 | NA | NO |
| p6.G3 | NA | M | 2018 | DEU | Det-2 | 48.556255 | 9.135424 | 0.374359 | 12.0216 | NA | NO |
| p7.F2 | NA | M | 2018 | DEU | Eyach | 48.446111 | 8.781611 | 0.420044 | 6.6193 | NA | NO |
| p7.G11 | NA | M | 2018 | DEU | Eyach | 48.446111 | 8.781611 | 0.922476 | 63.4571 | NA | YES |
| p8.B3 | NA | M | 2018 | DEU | Eyach | 48.446111 | 8.781611 | 0.999766 | 72.1507 | NA | YES |
| p8.B9 | NA | M | 2018 | DEU | Eyach | 48.446111 | 8.781611 | 0.921803 | 46.3749 | NA | YES |
| p8.C7 | NA | M | 2018 | DEU | Eyach | 48.446111 | 8.781611 | 0.933317 | 51.4209 | NA | YES |
| p8.D11 | NA | M | 2018 | DEU | Eyach | 48.446111 | 8.781611 | 0.432979 | 9.92024 | NA | NO |
| p8.E4 | NA | M | 2018 | DEU | Eyach | 48.446111 | 8.781611 | 0.917628 | 39.6076 | NA | YES |
| p8.G10 | NA | M | 2018 | DEU | Det-2 | 48.556255 | 9.135424 | 0.460138 | 12.253 | NA | NO |
| p8.G2 | NA | M | 2018 | DEU | Det-2 | 48.556255 | 9.135424 | 0.450816 | 8.26752 | NA | NO |
| p8.H7 | NA | M | 2018 | DEU | Det-2 | 48.556255 | 9.135424 | 0.899835 | 67.821 | NA | YES |
| p9.C4 | NA | M | 2018 | DEU | Eyach | 48.446111 | 8.781611 | 0.946827 | 40.897 | NA | NO |
| p9.H10 | NA | M | 2018 | DEU | Eyach | 48.446111 | 8.781611 | 0.469617 | 11.8808 | NA | NO |
| BAV2572 | NA | M | 2013 | USA | Virginia | NA | NA | NA | NA | NA | NO |
| CDRTc14 | NA | M | 2013 | AUT | Steinberg-Dörfel | 47.4N | 16.47E | NA | NA | NA | NO |
| ICMP8820 | NA | M | 2010 | NZL | NA | NA | NA | NA | NA | NA | NO |
| ICMP3272 | NA | M | 1975 | NZL | Riverhead | NA | NA | NA | NA | NA | NO |
| CFBP1590 | NA | M | 1974 | FRA | Vaucluse | NA | NA | NA | NA | NA | NO |
| DSM6694 | NA | M | 1930 | CHE | NA | NA | NA | NA | NA | NA | NO |
| CH409132 | NA | M | 2013 | USA | Boston, Massachusetts | 42.22N | 71.7W | NA | NA | NA | NO |
| LMCA8460 | NA | M | 2008 | USA | Lake Michigan College | 42.090114N | 86.393408W | NA | NA | NA | NO |
| SV1779 | NA | M | 2017 | USA | Scott Valley, California | NA | NA | NA | NA | NA | NO |
| T157 | NA | M | 2019 | USA | Tulelake, California | NA | NA | NA | NA | NA | NO |

**Table S4:** Key Resources Table.

| REAGENT or RESOURCE | SOURCE | IDENTIFIER |
| --- | --- | --- |
| **Deposited Data** |  |  |
| 1524 *Pseudomonas* genomes | Karasov et al. 2018 | ENA: PRJEB24450 |
| **Experimental Models: Organisms/Strains** |  |  |
| 1524 Modern *Pseudomonas* strains | Karasov et al. 2018 | NA |
| 34 Historic strains |  |  |
| 10 NCBI genomes | NCBI | In Table S3 (Sample) |
| p25.A12 ΔTFA<br>p25.A12ΔTFA +TFA | This study | NA |
| **Software and Algorithms** |  |  |
| VIBRANT v1.2.1 | Kieft et al., 2020 | https://github.com/AnantharamanLab/VIBRANT |
| Mash v2.2.2 | Ondov et al., 2016 | https://github.com/marbl/Ma |
|  |  | <u>sh</u> |
| panX | Ding et al., 2018 | <u>https://github.com/neherlab/pan-genome-analysis</u> |
| pankmer | Aylward et al., 2023 | <u>https://pypi.org/project/pankmer/</u> |

**Table S5.** De-novo assembly contigs statistics from reads mapped to the Tailocin region.

| MAETRIC\SAMPLE | HB0737 | HB0766 | HB0814 |
| --- | --- | --- | --- |
| **Total_n** | 10 | 9 | 14 |
| **Total_seq_bp** | 23769 | 21573 | 21574 |
| **Avg._seq_bp** | 2376.9 | 2397 | 1541 |
| **Median_seq_bp** | 2143 | 1665 | 1275.5 |
| **N_50_bp** | 2858 | 3405 | 1705 |
| **Min_seq_bp** | 583 | 494 | 415 |
| **Max_seq_bp** | 5543 | 7935 | 3337 |

## Notes

### Competing Interest Statement

The authors have declared no competing interest.

### Summary of Updates

We have substantially expanded the experiments and analyses of the manuscript to identify the tailocin receptor and determine the coevolution of the receptor in natural populations of Pseudomonas. To this aim, we bring to our team experts in glycobiology from the Georgia Carbohydrate Research Center. The set of experiments includes: 1. TnSeq Mutagenesis screen to identify important LPS receptors(Figure4A) 2. Mass-spectrometry of LPScomposition (Figure 4C,Table S2) and further carbohydrate analysis (Figure 4B). 3.Plant infection assays to test the in vivo effect of tailocins. With these new experiments we have determined: 1. That components of the lipopolysaccharide(LPS)O-chain are the receptor for the tailocin in the target cell. 2.The LPS residues,in concert with the tailocin tail fibers, are co-evolving in the Pseudomonas populations. 3. The Pseudomonas population that colonizes A. thaliana is polymorphic in the production of rhamnose residues in the LPS and mutation of these residues confers resistance. 4.The tailocin is naturally induced in plant infections and shows killing activity of neighboring bacteria in plant infections. Our co-infection results reveal that a wild type tailocin-encoding strain can completely suppress a competing strain in a plant infection.

